# Relation of CMV and brain atrophy to trajectories of immunosenescence in diverse populations

**DOI:** 10.1101/2024.10.07.614568

**Authors:** Hanane Touil, Tain Luquez, Natacha Comandante-Lou, Annie J Lee, Masashi Fujita, Christian Habeck, Alex Kroshilina, Everardo Hegewisch-Solloa, Julie McInvale, Leah Zuroff, Stephane Isnard, Elisabeth Walker, Lili Zhang, Jean-Pierre Routy, Ya Zhang, Emily M. Mace, Luisa Klotz, Heinz Wiendl, Zongqi Xia, Amit Bar-Or, Vilas Menon, Yaakov Stern, Philip L. De Jager

## Abstract

Immunosenescence (ISC), the aging of the immune system, has largely been studied in populations of European descent. Here, circulating immune cell cytometric data from African-American, Hispanic, and non-Hispanic White participants were generated. Known and novel age effects were identified using either a meta-analysis approach or a parallel genetic approach. Most results are consistent across the three populations, but some cell populations display evidence of heterogeneity, such as a PD-L1^+^CD56^+^ NK cell subset. The study estimated “Immunological Age” (IA) during physiologic aging. While we found no relation of IA to Multiple Sclerosis, IA is associated with entorhinal cortex atrophy, a presymptomatic feature of Alzheimer’s disease, linking neurodegeneration and peripheral immunity. ISC trajectories were also inferred, highlighting age, CMV status, and genetic ancestry as key influences. Our assessment offers reference ISC trajectories for personalization of assessments of immune function over the life course in diverse populations.

## Introduction

Aging is a physiological process that affects the entire organism, including the immune system. Immunosenescence (ISC) is a term used to describe changes in the immune system that occur with advancing age. ISC is characterized by a slow decline of immune competence, decreased responses to vaccination, and increased vulnerability to infectious diseases and neoplasia. It is thought to be a result of age-associated changes in cell functions as well as the composition of the innate and the adaptive immune system. Some changes like thymic involution occur early in adult life while most changes begin to accelerate in the sixth decade of life. ISC is often accompanied by inflammaging, a low-grade inflammation characterized by enhanced levels of peripheral pro-inflammatory cytokines including IL-6 and TNF-α ^1^. Several factors may impact the process of immune senescence, including environmental factors, previous viral infections ^2,3^, and possibly genetic variation across human individuals ^4^. However, our understanding of the extent of these changes and of factors that help some aged individuals maintain high immune performance while others become fragile, remains incomplete.

Recent studies using cytometric profiling in large numbers of human participants have begun to robustly outline the cell populations in the peripheral immune system that change in frequency with advancing age, specifically in the T cell compartment – such as reduced frequencies of naïve T cells, increased frequencies of terminally differentiated memory T cells, and a reduced repertoire diversity ^3,5,6^. Other flow cytometry-based studies have also noted age-associated changes in B cells and natural killer (NK) cells ^7–9^ ^10^ ^11,12^. Evaluation of the peripheral blood mononuclear cell (PBMC) transcriptome in centenarians and atherosclerosis patients has also led to the identification of age-related genes ^13^ ^14^ ^15^.

While a limited number of studies characterizing large cohorts are available, most studies have small numbers of participants and focus on populations of European ancestry. It remains unclear whether cellular and functional changes related to ISC are comparable among populations of European ancestry and under-studied populations such as African American and Hispanic populations. Thus, further characterization of ISC that considers sources of interindividual variation in immune traits over the lifespan will help inform the management of clinical care for our increasingly diverse older population. In addition, such studies have the potential to inform disease-related investigations by facilitating the distinction of early disease-associated immune alterations from those related to natural aging. One example is multiple sclerosis (MS), an inflammatory disease of the central nervous system in which the frequency of acute inflammatory relapses wanes with advancing age. It remains unclear how the cumulative inflammatory events during the life course of MS patients overlap with trajectories of physiologic ISC ^17–19^.

Here, we deployed a structured cytometric evaluation of a diverse set of individuals (n=222) to evaluate the immune landscape of peripheral blood over the adult lifespan in ethnically diverse donors aged between 25 and 88 years old. We deploy standard and advanced analysis approaches to these multiparametric immunophenotyping data to shed light on age-associated immune cell subsets, propose trajectories of ISC, identify factors that relate to these changes, and assess their relevance to brain aging and MS as a representative inflammatory disease.

## Results

### Characteristics of participants and analytical approach

We characterized 222 participants without chronic inflammatory diseases and 20 untreated multiple sclerosis (MS) patients. The data from the latter individuals were generated in an integrated fashion with the reference data to provide an early contrast with an inflammatory disease (see **Supplementary Table 1** for demographic characteristics). However, the MS participants are only used at the end of our pre-planned analyses; they are not used in our evaluation of the reference individuals who do not have an inflammatory disease. The reference individuals come from two longitudinal studies (demographic characteristics in **Supplementary Table 1**): (1) the Reference Ability Neural Network (RANN) study that includes diverse participants sampled from six decades of life (n=205, 26-84 years old) to study cognitive reserve ^20^ and (2) the Offspring Study (Offspring) of cognitive aging in diverse populations (n=20, 29-88 years old) (demographic characteristics in **Supplementary Table 1**). RANN Participants shared a self-reported ancestry (**Supplementary Table 1**), and genotype data were collected from the RANN and Offspring participants. PBMC were isolated and cryopreserved for both studies using strict standard operating procedures (SOPs) developed by the Center for Translational and Computational Neuroimmunology core at Columbia university (**Supplementary Methods**).

To characterize the innate and adaptive peripheral immune system with advancing age, we designed two cytometry-based immunophenotyping panels: (1) a PBMC panel (15 markers) to capture all major peripheral immune cell types and (2) a T cell panel (12 markers) to capture higher-resolution T cell phenotypes (**Supplementary Table 2**). To minimize the effect of technical variation introduced by sample batches in the experimental pipeline, two team members completed discrete tasks in parallel in each batch, and each batch contained 8 samples. These samples were preselected to ensure that each batch had samples from each sex, each parent study, each self-reported ancestry, and representatives across the age distribution (see **Methods**). Batch was accounted across all analyses assessing associations of immune-cell frequencies with age using a linear mixed-effects model with sex as a fixed effect and batch as a random intercept (**Table 1, Supplementary Table 3**).

**Table 1:** Association between immune cell subsets and chronologic age. Statistical analysis summarizing association between immune cell frequencies and chronologic age considering a meta-analysis of three population groups based on self-reported ethnicities: African American (AA), Hispanics and non-Hispanic whites (NHW). A joint analysis based on genetic principal components (PC), and considering the impact of CMV seropositivity. The results of the three individual analyses for AA, H and NHW participants are presented in **Supplementary Table** 3. Statistically significant correlations include p<0.05 and FDR<0.005. Number of participants used for each type of analysis: Self-reported ethnicities analysis (n=185), joint analysis using genetic principal components (PC) (n=222), joint analysis using PC and CMV as covariates (n=201). Glossary: CM: CD27^+^ CD45RA^-^ Central Memory T cells, EM: CD27^-^ CD45RA^-^ T cells, Naïve: CD27^+^ CD45RA^+^ Naïve T cells, TEMRA: CD27^-^ D45RA^+^ T effector memory cells.

|  | Fixed effects meta-analysis of three population groups based on self-reported ethnicities |  |  |  |  |  | Joint-analysis using genetic Principal Component (PC) |  |  |  |  |  |  |  |  |  |
| --- | --- | --- | --- | --- | --- | --- | --- | --- | --- | --- | --- | --- | --- | --- | --- | --- |
|  |  |  |  |  |  |  | Joint analysis using genetic PC as covariates |  |  |  |  | Joint analysis using genetic PC & CMV as covariates |  |  |  |  |
| Immune cell | n | b | se | p | fdr | I<sup>2</sup> | n | b | se | p | fdr | n | b | se | p | fdr |
| CD8+CD56+ NK T cells | 185 | 0.003 | 0.002 | 0.082 | 0.164 | 50.200 | 222 | 0.003 | 0.001 | 0.002 | **0.007** | 201 | 0.001 | 0.001 | 0.179 | 0.274 |
| CD56+CD16+ NK cells | 185 | 0.008 | 0.004 | 0.048 | 0.128 | 23.019 | 221 | 0.006 | 0.003 | 0.061 | 0.106 | 201 | 0.002 | 0.003 | 0.506 | 0.626 |
| CD56+CD16+ PD-L1+ NK cells | 164 | -0.001 | 0.002 | 0.768 | 0.981 | 0 | 221 | -0.003 | 0.003 | 0.353 | 0.437 | 201 | -0.001 | 0.003 | 0.708 | 0.782 |
| CD56+PD-L1+ NK cells | 185 | 0.004 | 0.002 | 0.050 | 0.128 | 16.650 | 222 | 0.004 | 0.001 | 0.004 | **0.012** | 201 | 0.002 | 0.002 | 0.156 | 0.271 |
| **T cells** |  |  |  |  |  |  |  |  |  |  |  |  |  |  |  |  |
| CD3+ T cells | 185 | -0.1045 | 0.0514 | 0.0422 | 0.1284 | 0 | 222 | -0.122 | 0.046 | 0.009 | **0.023** | 201 | -0.152 | 0.050 | 0.003 | **0.017** |
| CD4+ T cells | 185 | -1.0863 | 23.1225 | 0.9625 | 0.9811 | 72.90 | 222 | 13.907 | 10.024 | 0.167 | 0.271 | 201 | 25.137 | 10.893 | 0.022 | 0.064 |
| CM CD4+ T cells | 185 | -0.0724 | 0.0668 | 0.2783 | 0.4522 | 0.02 | 222 | -0.045 | 0.059 | 0.445 | 0.526 | 201 | -0.076 | 0.066 | 0.247 | 0.356 |
| EM CD4+ T cells | 185 | 0.0095 | 0.0027 | 0.0005 | **0.0027** | 0 | 222 | 0.007 | 0.002 | 0.003 | **0.011** | 201 | 0.004 | 0.002 | 0.154 | 0.271 |
| Naïve CD4+ T cells | 185 | -0.0933 | 0.0803 | 0.2454 | 0.4254 | 0 | 222 | -0.088 | 0.070 | 0.209 | 0.302 | 201 | -0.002 | 0.075 | 0.981 | 0.981 |
| TEMRA CD4+ T cells | 185 | 0.0020 | 0.0006 | 0.0003 | **0.0027** | 0 | 222 | 0.002 | 0.001 | 0.003 | **0.010** | 201 | 0.001 | 0.001 | 0.057 | 0.136 |
| CD8+ T cells | 185 | -3.6E-05 | 1.5E-03 | 9.8E-01 | 9.8E-01 | 51.45 | 222 | -0.001 | 0.001 | 0.206 | 0.302 | 201 | -0.002 | 0.001 | 0.020 | 0.064 |
| CM CD8+ T cells | 185 | 6.9E-03 | 1.1E-02 | 5.3E-01 | 7.3E-01 | 49.26 | 222 | -0.001 | 0.006 | 0.885 | 0.885 | 201 | 0.005 | 0.007 | 0.419 | 0.573 |
| EM CD8+ T cells | 185 | 9.5E-03 | 2.7E-03 | 5.1E-04 | **2.7E-03** | 0.08 | 222 | 0.009 | 0.003 | 0.001 | **0.007** | 201 | 0.007 | 0.003 | 0.013 | 0.058 |
| Naïve CD8+ T cells | 185 | -0.84680 | 0.15425 | 4.03E-08 | **5.24E-07** | 64.37 | 222 | -0.664 | 0.069 | 2.40E-18 | **6.25E-17** | 201 | -0.582 | 0.071 | 6.01E-14 | **1.56E-12** |
| TEMRA CD8+ T cells | 185 | 0.02963 | 0.00420 | 1.71E-12 | **4.45E-11** | 0 | 222 | 0.026 | 0.004 | 1.99E-11 | **2.59E-10** | 201 | 0.020 | 0.004 | 5.30E-08 | **6.88E-07** |
| CD3+ CD8+PD1+ T cells | 184 | 0.000 | 0.002 | 0.878 | 0.981 | 29.264 | 221 | 0.001 | 0.001 | 0.564 | 0.638 | 200 | 0.000 | 0.001 | 0.930 | 0.967 |
| **B cells** |  |  |  |  |  |  |  |  |  |  |  |  |  |  |  |  |
| CD20+ B cells | 185 | -0.0064 | 0.0023 | 0.0046 | **0.0200** | 0 | 222 | -0.006 | 0.002 | 0.002 | **0.007** | 201 | -0.007 | 0.002 | 0.002 | **0.016** |
| Memory IgD+/-CD27+CD20+ B cells | 185 | -0.0016 | 0.0021 | 0.4470 | 0.6836 | 56.36 | 222 | -0.002 | 0.001 | 0.051 | 0.106 | 201 | -0.003 | 0.001 | 0.019 | 0.064 |
| Naïve IgD+CD20+ B cells | 185 | -1.2664 | 7.6363 | 0.8683 | 0.9811 | 47.24 | 222 | 0.777 | 4.259 | 0.855 | 0.885 | 201 | 3.565 | 4.737 | 0.453 | 0.589 |
| CD11c+ CD20+ B cells | 185 | 0.0088 | 0.0049 | 0.0698 | 0.1513 | 68.61 | 222 | 0.007 | 0.002 | 0.001 | **0.007** | 201 | 0.006 | 0.002 | 0.011 | 0.058 |
| Memory IgD-CD27+ CD20+ B cells | 185 | -0.0008 | 0.0012 | 0.5112 | 0.7311 | 12.07 | 222 | 0.000 | 0.001 | 0.624 | 0.676 | 201 | -0.001 | 0.001 | 0.175 | 0.274 |
| **Monocytes** |  |  |  |  |  |  |  |  |  |  |  |  |  |  |  |  |
| Classic CD14+CD16- Monocytes | 185 | 0.0011 | 0.0087 | 0.8984 | 0.9811 | 50.52 | 221 | -0.005 | 0.005 | 0.249 | 0.340 | 201 | -0.003 | 0.005 | 0.595 | 0.704 |
| Intermediate CD14highCD16+ Monocytes | 185 | 0.0006 | 0.0003 | 0.0543 | 0.1284 | 0 | 221 | 0.001 | 0.000 | 0.061 | 0.106 | 201 | 0.001 | 0.000 | 0.031 | 0.081 |
| Non-classical CD14low CD16high Monocytes | 185 | 0.0042 | 0.0018 | 0.0208 | 0.0774 | 51.78 | 221 | 0.003 | 0.001 | 0.010 | **0.023** | 201 | 0.002 | 0.001 | 0.127 | 0.254 |
| CD11cHigh CD14High Monocytes | 185 | -0.0003 | 0.0073 | 0.9668 | 0.9811 | 46.90 | 222 | -0.004 | 0.004 | 0.290 | 0.377 | 201 | -0.002 | 0.004 | 0.722 | 0.782 |
| Live cells | 185 | -0.0056 | 0.0042 | 0.1859 | 0.3452 | 0 | 222 | -0.007 | 0.004 | 0.059 | 0.106 | 201 | -0.007 | 0.004 | 0.094 | 0.205 |

Given that cytomegalovirus (CMV) seropositivity status is known to affect the frequency of certain immune cell subsets, including T effector memory cells and natural killer (NK) cells ^3^, we assessed CMV serology and measured anti-CMV IgG titers using available serum samples from the RANN cohort (n=201) collected from the same blood draw as the PBMC (**Supplementary Table 4**). Functional magnetic resonance imaging (MRI) were collected for RANN participants as part of the parent protocol ^21^ (n=185 have both cytometric and MRI data).

Flow cytometry data were analyzed using two parallel approaches: (1) a standard manual segmentation approach using FlowJo 10.7 (Becton Dickinson BD, NJ) software and sequences of two-dimensional gates which captured 28 manually curated distinct immune cell subtypes across the two panels, and (2) an automated segmentation approach using PhenoGraph which uncovered 35 (PBMC panel) and 32 (T cell panel) cell subtypes for analysis (see **Methods**).

### Association between chronological age and manually-segmented cell subsets among individuals of different ancestries

We first sought to assess the impact of chronological age on the frequency of defined immune cell subsets (see **Methods**). Since we have three major groups of participants based on their self-reported race and ethnicity – non-Hispanic African-American (AA), Hispanic (H), and non-Hispanic White (NHW) participants – we analyzed our data in each of the three groups separately and combined the results using a random-effects meta-analysis approach (up to 185 participants) (**Table 1, Supplementary Table 3**) for one of our two primary analyses. A false discovery rate (FDR) <0.05 was used to account for the testing of multiple hypotheses. We note that a small number of self-reported Asian, Pacific Islander, biracial and “Other” participants (**Supplementary Table 1**) were not included in the three ethnic group meta-analysis due to their limited number.

Published studies have reported changes in immune cell populations with advancing age ^3,4,22,23^. Thus, we first sought to validate age-related changes and confirmed statistically significant effects of age on the frequency of the following immune cell subsets (FDR<0.05 in the meta-analysis) (**Table 1**): (1) an increase in T effector memory (TEM) cells and TEM cells that co-express CD45RA (TEMRA) (both CD8^+^ T cell subtypes), **Figure 2B, D** and **supplementary Figure 1 E)**, (2) a decrease in naïve CD8 T cells (**Figure 2 B and C**), (3) an increase in TEM and TEMRA CD4^+^ cells (**Figure 2 E, G** and **Supplementary Figure 1 G)**, and (4) a decrease in CD20^+^ B cells (**Supplementary Figure 1 C** and **D**). There have been conflicting reports about CD11c^+^ B cells as a hallmark of immune-aging ^10,24^ perhaps due to the heterogeneity of this subset co-expressing CD21, CD38, CD11b and CD11c; here, our data suggest that CD11c^+^CD20^+^ B cells are significantly increased in frequency with advancing age but only among AA participants (**Figure 2 H, I & J, Supplementary Table 3**); this result is not significant in the meta-analysis.

**Figure 1:**
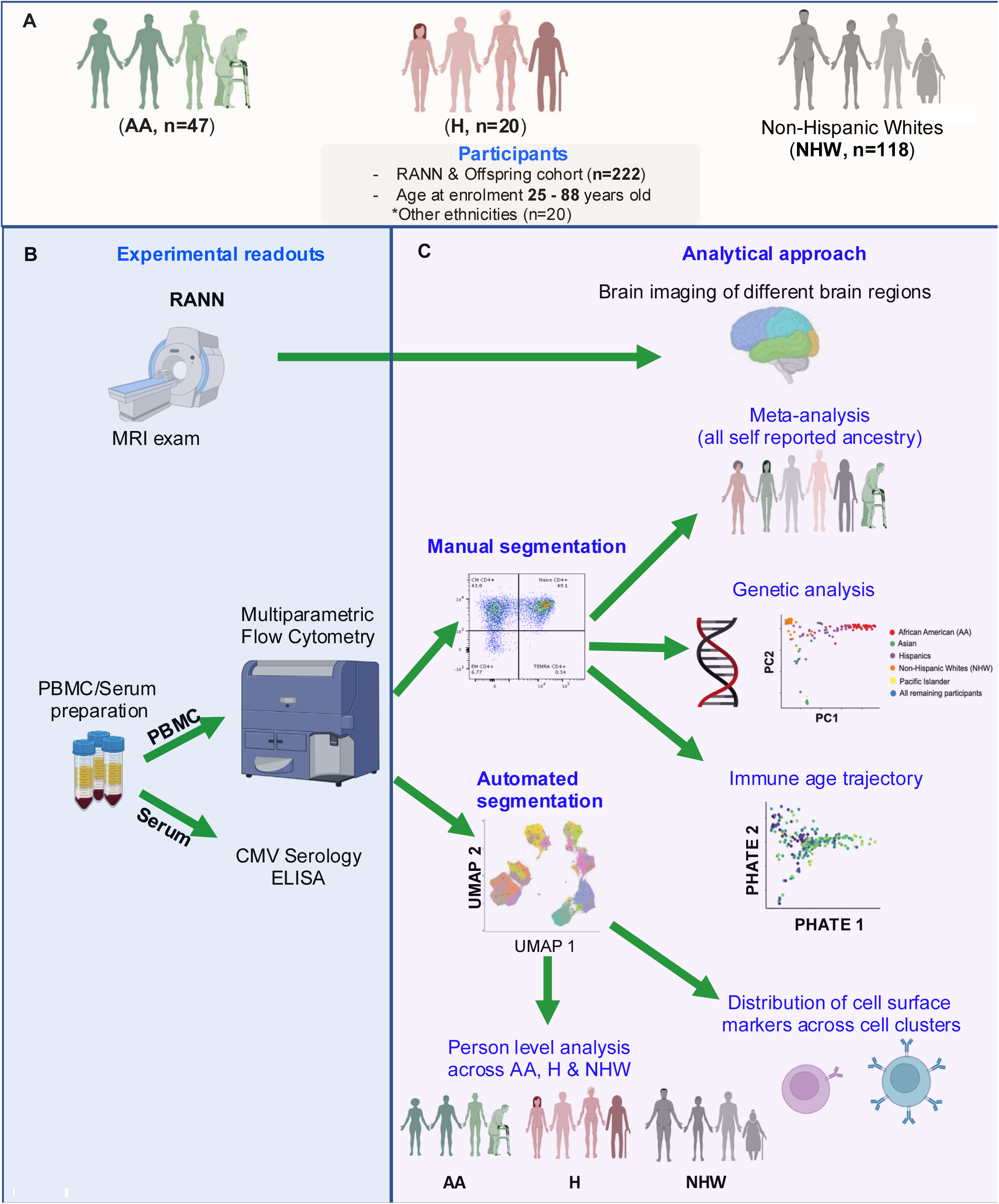
Study Design. (**A**) Represents a description of cohort considered to characteries trajectories of immune-aging in this study. (**B**) Detailed experimental readouts ran on the study participants. (**C**) Analytical approach used for Flow Cytometry data, included the manual segmentation using FlowJo, and the automated segmentation using PhenoGraph.

**Figure 2:**
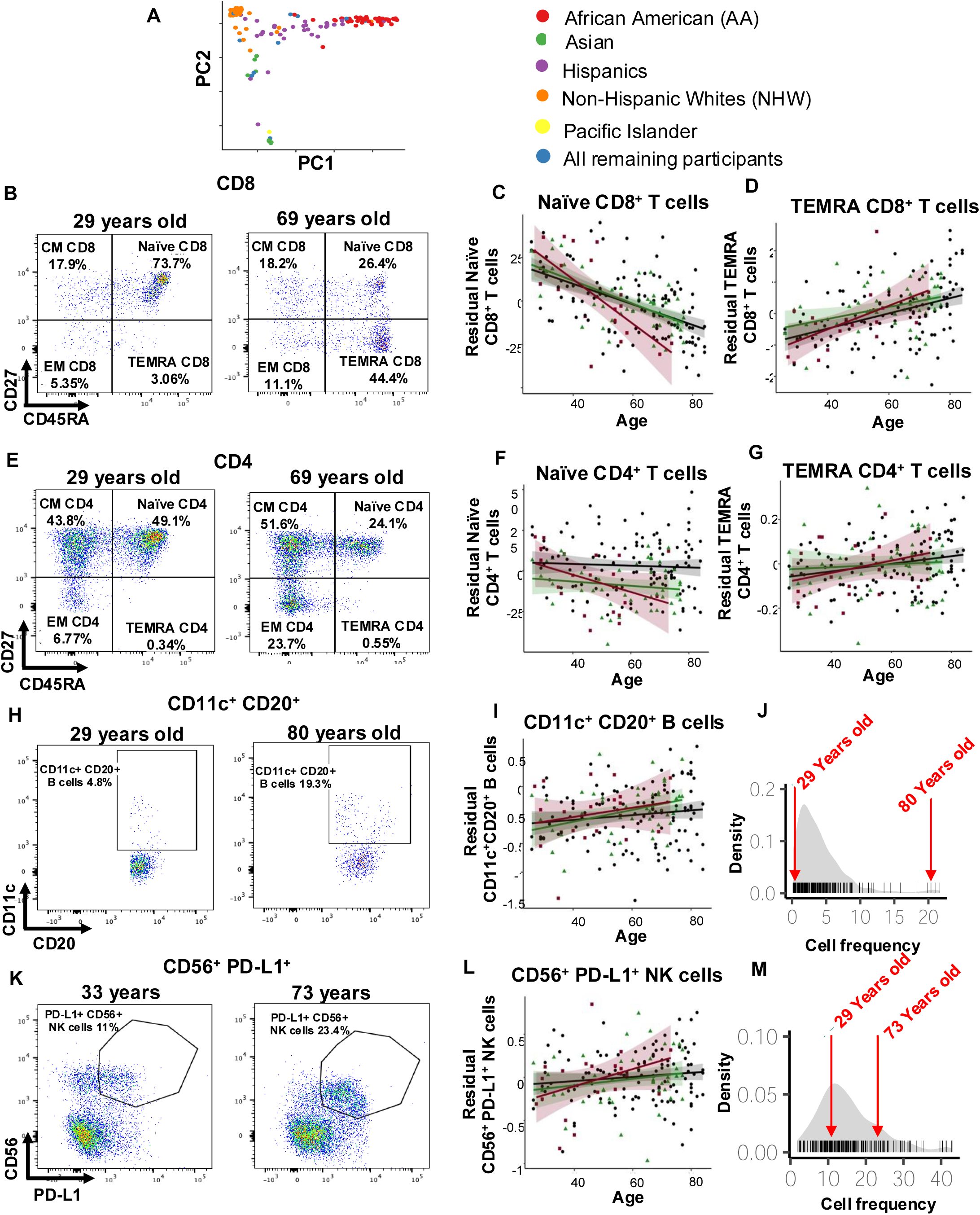
Illustration of known and novel effects of age on cell population frequency between chronological age and manually segmented cell subsets among individuals of different self-reported ancestries. **(A)** Representation of broad ancestry variation based on principal components (PC) derived from our study participants and obtained from genome wide data**. (B** to **M**) Cryopreserved human peripheral mononuclear cells (PBMC) were thawed and stained with cell surface markers, and further analyzed with Flow Cytometry. (**B**) Representative Flow Cytometry plots of a 29 and a 69-year-old individual highlighting the different CD8 T cell subsets including Central Memory (CM), Naïve, Effector Memory (EM) and T effector Memory cells co-expressing CD45RA (TEMRA) T cells. (**C**) Plots demonstrating the visual representation of the relationship between frequencies of Naïve and (**D**) TEMRA CD8 T cells with chronological age using distinct linear regression curves shown for African American (AA) (green curves), Hispanics (burgundy curves), and non-Hispanic whites (NHW) (black curves). (**E**) Representative Flow Cytometry plots of a 29 and a 69-year-old individual highlighting the different CD4 T cell subsets including Central Memory (CM), Naïve, Effector Memory (EM) and T effector Memory cells co-expressing CD45RA (TEMRA) T cells. (**F**) Plots demonstrating the visual representation of the relationship between frequencies of Naïve and (**G**) TEMRA CD4 T cells with chronological age using distinct linear regression curves shown for African American (AA) (green curves), Hispanics (burgundy curves), and non-Hispanic whites (NHW) (black curves). (**H**) Representative Flow Cytometry plots of CD11c^+^ CD20^+^ B cells derived from a 29 and a 78-year-old individual. B cells were gated on CD3^-^ cells, and further gated on CD20^+^ B cells co-expressing CD11c. (**I**) Plots demonstrating the visual representation of the relationship between frequencies of CD11c^+^ CD20^+^ B cells and chronological age using distinct linear regression curves shown for Hispanics (burgundy curves), non-Hispanic whites (NHW) (black curves) and African American (AA) (green curves). (**J**) Distribution curve of CD11c^+^CD20^+^ B cell frequencies showing a representation of a 29 and an 89-year-old participant. (**K**) Representative Flow Cytometry plots of CD56^+^ PD-L1^+^ NK cells derived from a 29 and a 73-year-old individual. NK cells were gated on CD3^-^ cells, and further gated on CD56 co-expressing PD-L1. (**L**) Plots demonstrating the visual representation of the relationship between frequencies of CD56^+^ PD-L1^+^ NK cells and chronological age using distinct linear regression curves shown for African American (AA) (green curves), Hispanics (burgundy curves), non-Hispanic whites (NHW) (black curves). (**M**) Distribution curve of CD56c^+^PD-L1^+^ NK cell frequencies showing a representation of a 29 and a 73-year-old participant. Statistical analysis is detailed in **Table 1**.

To assess whether the effect of age is different for certain cell types among the three self-reported population groups, we calculated a measure of heterogeneity – the I^2^ statistic (**Table 1**) - in the effect sizes of the three populations included in the meta-analysis. Interestingly, the cell types fell into two separate groups (**Table 1**): (1) cell populations (such as B cells (**Supplementary Figure 1 D**) as well as CD4+ and CD8+ TEMRA (**Figure 2 D and G)** and CD4^+^ TEM cells **(Supplementary Figure 1 G**) which have an I^2^=0, meaning that there are consistent effects of age in all three populations, and (2) cell populations that have either moderate (I^2^>30) or substantial (I^2^>50) evidence of heterogeneity in the effect of age, such as Naïve CD8^+^ T cells (**Figure 1 C, Supplementary Table 3**) which change more profoundly in individuals who are not of Hispanic ancestry (**Table 1, Supplementary Table 3**). This is also seen with the CD11c^+^CD20^+^ B cell subtype (**Figure 2 H** and **I, Table 1**) for which the increase in frequency with age is much greater in AA participants and non-significant among NHW participants, despite a much larger sample size. Another example is the CD56^+^CD8^+^ NKT cell subtype (**Supplementary Figure 1 B, Supplementary Table 3)**: here, the effect of age is greater among NHW than in the other two groups.

To complement the meta-analysis approach that utilizes self-reported ancestry, we deployed a second primary analysis in which we corrected our analysis of age effects using estimates of genetic ancestry based on genome-wide genotype data (see **Methods**, **Table 1**). The advantage of this approach is that we were able to use all participants with available genome-wide genotype data (222 participants), which included those individuals who had reported their ancestry as “other”. The top three principal components (PC) from the genome-wide data were included as covariates in the linear regression analysis to account for broad effects of ancestry. The genotype-corrected analysis discovered the same associations as the meta-analysis of self-reported ancestries and identified five additional cell populations with significant changes (FDR<0.05) (**Table 1** and **Supplementary Table 3)**: the CD11c^+^CD20^+^ B cells and CD56^+^CD8^+^ NK cell subtypes discussed above which have a suggestive association in the meta-analysis as well as a decrease in CD3+ T cells, an increase in CD56^+^PD-L1^+^ NK cells and an increase in non-classical CD14^low^CD16^high^ monocytes with advancing age. The latter observation replicates a prior report associating an increase in this cell population with advancing age ^25^, but the CD56^+^PD-L1^+^ NK cell association has not been reported previously. Thus, while we used two very different analytic methods to account for race and ethnicity in our participants, results are relatively robust to either a self-reported ancestry or a genotype-based approach **(Table 1, Supplementary Table 3 and 6,** respectively). It does appear that the genotype-based approach may be better powered since it uncovered five additional age-affected cell populations in a sample that is slightly larger than that used in the meta-analysis. Indeed, from the meta-analysis results, we see that these two populations (CD56^+^PD-L1^+^ NK cells and CD14^low^CD16^high^ monocytes) were trending in the same direction but did not reach significance.

Previous studies have noted that some age-related changes in cell populations may in fact be related to exposure to cytomegalovirus (CMV) ^3^. We therefore repeated the genotype-based analysis adding a covariate for exposure to CMV. **Table 1** shows the results: most of the age associations are strongly attenuated when we account for the exposure to CMV, although four associations remain significant. Since we do not have longitudinal samples, we cannot resolve the causal relationship between aging and CMV exposure on PBMC subtype frequencies at this time.

### Association between chronological age and cell subtypes from automated segmentation of the PBMC and T cell cytometric profiles

Given that the richness of our high-dimensional single cell cytometric data is not fully leveraged by traditional, manual segmentation, we implemented automated segmentation that considers all dimensions of information simultaneously using PhenoGraph ^26^ (see **Methods**). Overall, we identified 35 distinct cell clusters within the PBMC dataset (**Figure 3 A**) and 32 cell clusters within the T cell dataset (**Supplementary Figure 2 A**). The T cell population encompasses 16 distinct subclusters, while myeloid cells, B cells and NK cells include between 4-6 distinct cell subclusters. The multidimensional analysis validated our manually segmented findings including changes within the naïve and memory CD8^+^ T cells (**Figure 2 C** and **D**). However, PhenoGraph expands the number of sub-clusters being considered, especially among NK and T cell populations (**Figure 3 B** and **supplementary Figure 2 B**).

**Figure 3:**
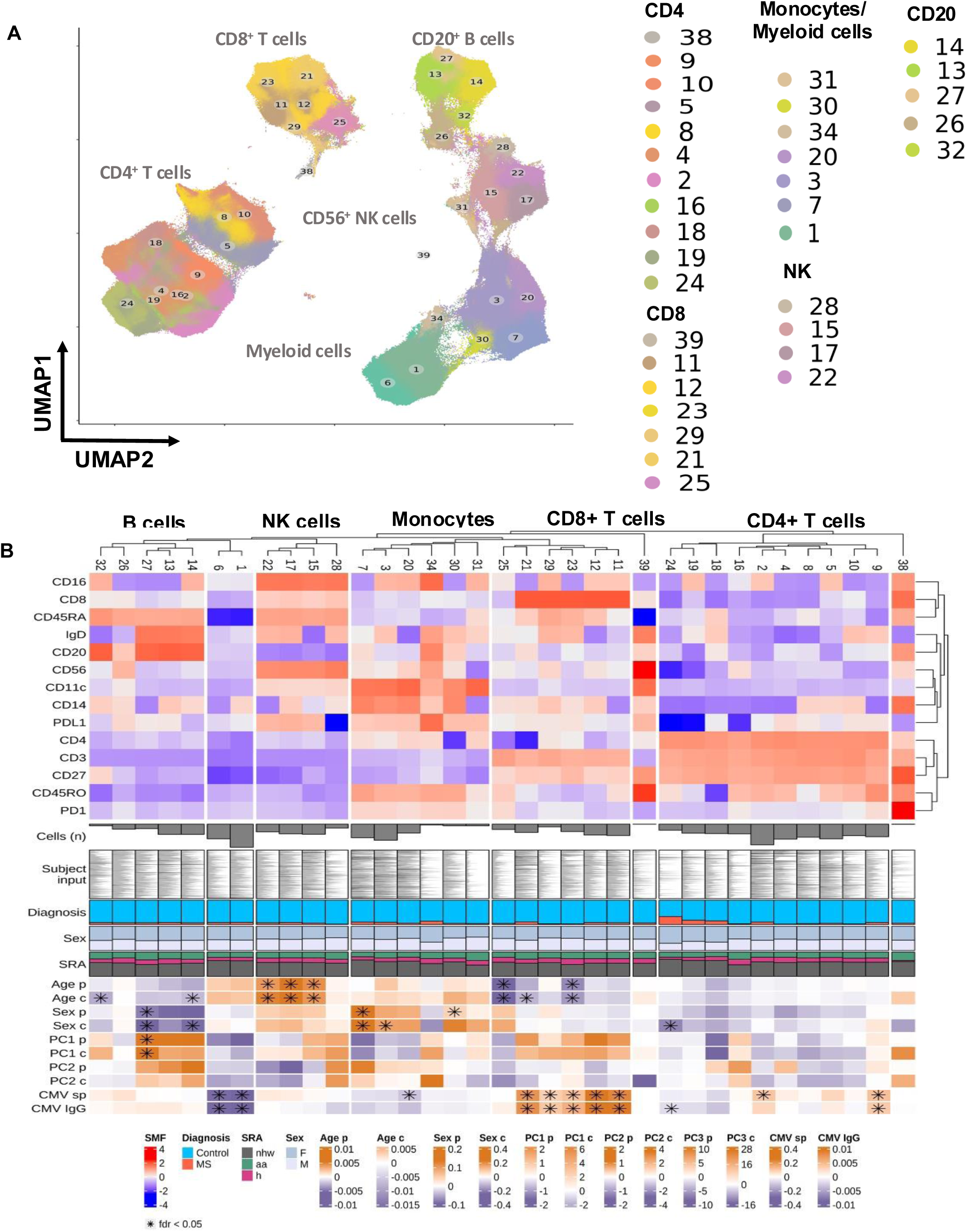
Automated cell clustering reveals novel cell clusters associated with chronological age among individuals of different self-reported ancestries. (**A**) Uniform manifold approximation and projection (UMPA) visualization plot of 39 PBMC clusters based on the cell surface panel markers (CD3, CD4, CD8, CD11c, CD14, CD16, CD20, CD27, CD45RA, CD45RO, CD56, PD1 & PD-L1). (**B**) Heatmap depict fluorescence of each cell surface marker that help define the different immune cell subsets within 247 individuals (African American (AA), Hispanics, and non-Hispanic whites (NHW)), and 20 untreated RRMS patients. Fluorescence ranges from −4 (blue) to +4 (red). Cell surface markers are used are presented across the top y-axis. Cell cluster numbers representing distinct cell subsets are ordered on top of the heatmap. The number of cells (n cell) within each cell cluster is expressed in the x-axis. The proportion (prop) of each cell subset across each cell cluster is represented in the x-axis. The diagnosis of each study participant is represented in the x-axis, blue represents control participants and red represents untreated Relapsing Remitting Multiple Sclerosis patients (RRMS). The distribution of female (light blue) or male (purple) study participants is represented in the x-axis. The distribution of study participant’s self-reported ancestries (SRA) including African American (AA) (green), Hispanics (burgundy), non-Hispanic whites (NHW) (black) is represented in the x-axis. Heatmap depict statistical analysis of cell cluster association with: age at the participant level (Age p) FDR values ranging from 0.01 (orange) to −0.01 (purple), the cell level (Age c) FDR values ranging from 1.02 (orange) to 0.98 (purple); with sex at the participant level (Sex p) FDR values ranging from 0.2 (orange) to 0.6 (purple) and the cell level (Sex c) FDR values ranging from 1.4 (orange) to 0.6 (purple) and with genetic ancestry PC1 p, PC2 p and PC3 p at the participant level FDR values ranging from 3 (orange) to −2 (purple) and the cell level PC1 c, PC2 c and PC3 c FDR values ranging from 1000 (dark orange) to 200 (light orange). * Marks statistically significant associations with FDR <0.05.

To identify putative age associations after adjusting for genetic ancestry (using the approach described above in analyzing the manually segmented data), we used two approaches: 1) a linear mixed-effect model with the person-level proportions of each cell type and 2) a cell-level logistic mixed-effect model that leverages the information from cell-to-cell variation across all individuals (see **Methods**). The person-level analysis highlights three distinct PD-L1^+^ NK cell subsets (cluster 15, FDR=0.03; cluster 17, FDR=0.001; & cluster 22, FDR=0.002, all clusters within the PBMC panel) that increase with advancing age (**Figure 3 B, Supplementary Figure 1 I-K**). These three NK cell subsets are distinguished by differences in the relative expression of IgD and CD14 (**Figure 3 B**). Furthermore, we report a significant decrease in the frequency of CD3^+^CD45RO^+^PD-L1^+^ T cells (cluster 25 (PBMC panel), FDR=6.09 x 10^−5^) and CD8^+^CD45RA^+^IgD^+^CD27^high^ T cells (cluster 23 (PBMC panel), FDR=0.00013) (**Figure 3 B, Supplementary Table 7**). On the other hand, the cell-level analysis (**Figure 3 B**) uncovers the same associations and three additional ones: another CD8^+^ T cell (cluster 29, FDR=6.34 x 10^−9^ - T cell panel) expressing the senescence marker KLRG1 **(Supplementary Figure 2 B** and **E**) and two B cell populations IgD^+^ naïve B cells (cluster 14, FDR=0.03 - PBMC panel) and the CD27^+^ cluster 32, FDR=0.01 - PBMC panel) (**Figure 3 B** and **Supplementary Table 7**). Thus, the cell-level analysis may provide increased sensitivity in uncovering age-related effects. Overall, we replicated all established immune cell changes with age, except for the associations with CD8^+^ TEMRA and CD11c^+^ B cells as these cell types seem to be distributed across several clusters with the phonograph-based segmentation.

We next assessed to what extent the abundance of cell surface proteins within each cluster is impacted by advancing age using the same analysis strategy. We observed a reduced CD27 mean fluorescence intensity with age in five out of six CD8^+^ T cell clusters (cluster 11, 12, 21, 23 & 29 - PBMC panel) (**Supplementary Figure 3**). Two of these clusters (21 and 23), also show a significant reduction of CD45RA **Supplementary Figure 3**), while cluster 23 also shows an increased expression of PD-L1 and CD45RO over time. Since the frequency of these populations are also associated with CMV, the two may be related, consistent with a loss of CD8^+^ T cells with an early differentiation phenotype as the participant ages.

Overall, the automated segmentation not only recapitulates the manual segmentation results but also, by considering all multidimensional data simultaneously, leads to a more refined cell subtype characterization compared to the classic approach using a series of two-dimensional gates. Where differences arise between automated and manually segmented data (as in this study), it is important to highlight them to guide further evaluations that can refine cell subtype definitions and take advantage of different analytic approaches such as the cell-level that is more sensitive.

### Impact of sex, genetic ancestry, and CMV infection on trajectories of immune aging

To thoroughly evaluate the role of other pertinent variables, we performed analyses to assess the effect of sex, independent of age, and we found that in the PBMC panel naïve CD20^+^IgD^+^ B cells (cluster 27, FDR=0.03), CD45RA^+^ pre-B cells (cluster 14, FDR=0.01), myeloid cells (clusters 3, FDR=0.04 and cluster 7, FDR=0.006), and CM CD4 T cells (cluster 24, FDR=0.02) are influenced by sex (**Figure 3 B**). The T cell panel further resolved the T cell association with sex and uncovered an effect on naïve CD27^+^ CD45RA^+^ CD8^+^ subtype (cluster 24, FDR=0.01), and naive CD27^+^ CD45RA^+^ CD4^+^ T cells (cluster 6, FDR=0.03) (**Supplementary Figure 2 B, Supplementary Table 8**). On the other hand, the first principal component (PC1) derived from genome-wide genotype data - which captures most of the ancestry gradient between European and African ancestry (**Figure 1 A**) – was only associated with an increase in the frequency of PBMC cluster 27 (p=0.004) (an IgD^+^ naïve B cell subtype) that is also influenced by sex (**Figure 3 B).** The other genetic PC (PC 2) is not significantly associated with cytometric traits, so the effect of genetic ancestry on our cytometric data is relatively modest when compared to other factors such as earlier CMV exposure.

In fact, we find that CMV infection is associated with increased proportions of five CD8^+^ T cell-subsets (cluster 11, FDR=1.06 x10^−5^, cluster 12, FDR=4.93 x10^−6^, cluster 21, FDR=7.42×10^−7^, cluster 23, FDR=0.003, and cluster 29, FDR=0.007) and two CD4 T cell subsets (cluster 2, FDR=0.028 and cluster 9, FDR=0.001) (**Figure 3 B**). While the myeloid cell cluster 20 seems to be only significantly associated with CMV seropositivity (FDR=0.03, **Supplementary Table 7**) but not with the titer of these immunoglobulins (FDR=0.09, **Figure 3 B).** Additionally, CMV titers (**Supplementary Table 4**) - but not seropositivity - are associated with a reduction in a subset of naïve CD4^+^ T cells (cluster 14, FDR=0.03), and CD8^+^ T cells co-expressing KLRG1 (cluster 12, FDR=0.0001) (**Supplementary Figure 2 B**, and **Supplementary Table 8**). As noted above, many of these cell populations are also associated with age, and so ambiguity remains at this time in terms of whether the cell population changes are primarily due to an increased risk of CMV infection with advancing age or an acceleration of immune aging after infection.

### Age-associated PD-L1^+^ NK cells are mature and functionally distinct

Having uncovered an NK cell subset co-expressing CD56 and PD-L1 (**Figure 3 B**) that substantially increases in frequency with aging, we further explored the transcriptional and proteomic profile of this cell subtype by interrogating CITE-seq data generated from 12 of our participants. We segmented these data based on the 14 markers that are shared between the PBMC cytometric panel and the CITE-seq panel, gated out CD3^+^ T cells, CD14^+^ myeloid cells and CD19^+^ B cells and further selected 4 NK clusters (cluster 22, 17, 15 and 28) as the preferred resolution in an attempt to parallel our automated segmentation-based analyses (**Figure 4 A**) (see **Supplementary Methods**). Here, we find one NK subtype with a high expression of PD-L1 (NK cell cluster 13), and we compared this subtype to NK cell cluster 2 and NK cell cluster 22 (**Figure 4 A**) which express the lowest levels of PD-L1 and are used as reference populations.

**Figure 4:**
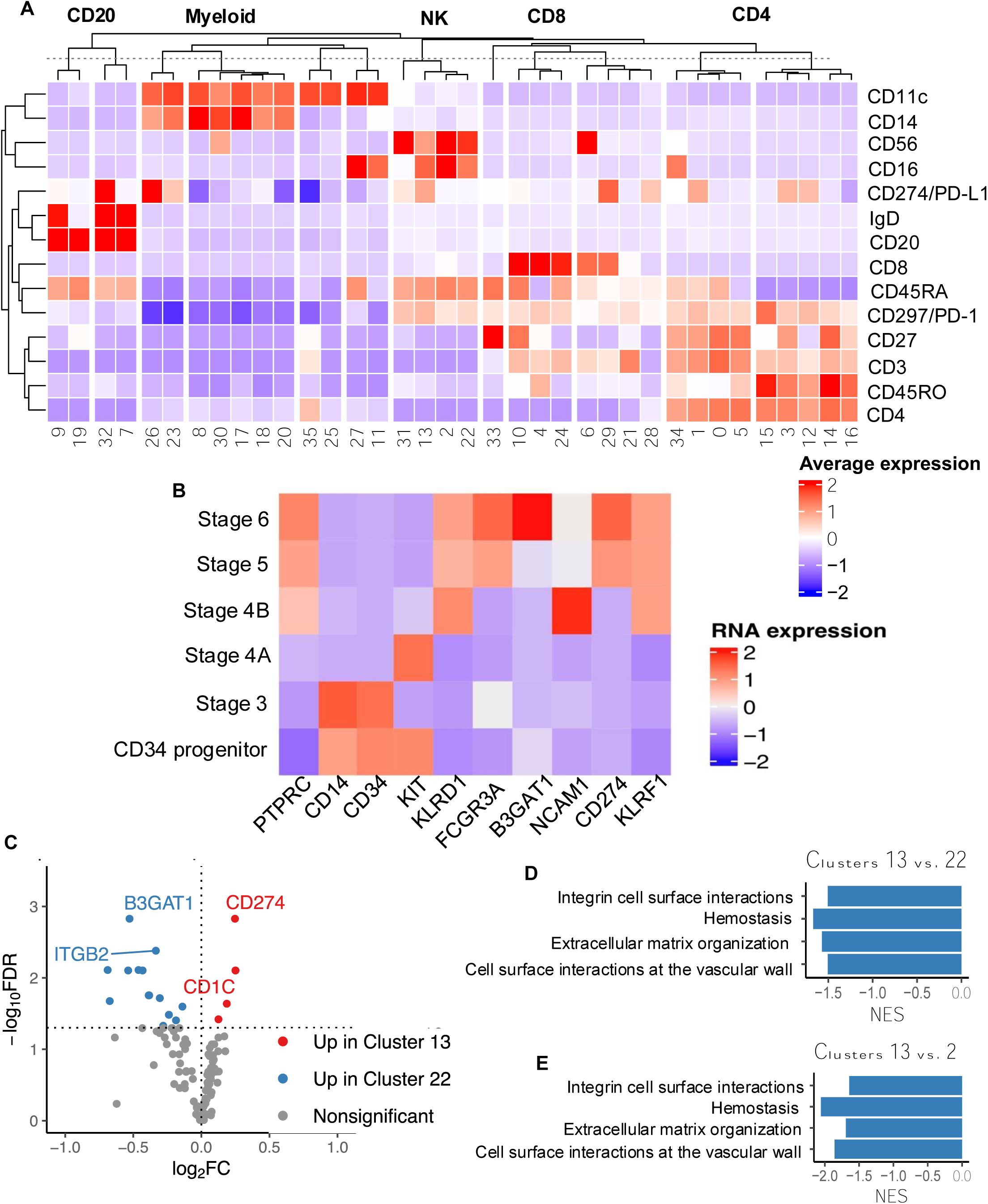
Characterization of the age associated CD56+ PD-L1+ NK cell subset. Cryopreserved human peripheral mononuclear cells (PBMCs) were thawed and stained with cell the CITE-Seq panel C (**A**) Heatmap depict protein expression of 14 markers out of 137 markers within the CITE-Seq panel that help define immune cell clusters identified using the automated flow cytometry analysis. (**B**) Heatmap depict gene expression of 9 genes within the different NK cell maturation stages, including distribution of CD247 coding for PD-L1 across the different maturation stages. Cells were first gated on live CD56^+^ NK cells, then gated on CD45+ and CD3^-^ lymphocytes. (**C**) Pairwise differential gene expression (DEG) comparing cluster 13 (co-expressing CD56 and PD-L1), with cluster 22 (that do not express PD-L1). Genes labeled in red are up-regulated in cluster 13 and down-regulated in cluster 22. Genes labeled in blue are up-regulated in cluster 22 and down-regulated in cluster 13. Pathway enrichment analysis using Reactome database (**D**) Bar plot representing normalized enrichment scores (NES) within pathways enriched in cluster 13 (CD56^+^ PD-L1^+^) compared to cluster 22 (CD56^+^PD-L1^-^), (**E**) Bar plot representing normalized enrichment scores (NES) within pathways enriched in cluster 13 (CD56^+^ PD-L1^+^) compared to cluster 2 (CD56^+^ PD-L1^-^).

Differential gene and protein expression in the CITE-seq data confirmed a higher expression of PD-L1 in cluster 13 relative to clusters 22 (**Figure 4 C**); CD1c which is expressed at varying levels by antigen presenting cells - is also more highly expressed in cluster 13 compared to cluster 22, but there is no significant difference of CD1c expression between cluster 13 and cluster 2 (**Supplementary Figure 5 H and Supplementary Table 9**). Hence, CD56^+^ PD-L1^+^ NK cells are a transcriptionally distinct population with some genes suggesting antigen presenting features, although further investigation is needed to confirm antigen presenting function. Gene set enrichment analysis using the Reactome database was performed comparing PD-L1^+^ NK cells (cluster 13) with PD-L1^-^ NK cells (clusters 22 and 2). We report a substantial enrichment of pathways involved in integrin mediated cell surface interactions (p=0.02), in cell surface interaction at the vascular wall (p=0.0018), in cell homeostasis (p=5.82 x10^−6^), and in extracellular matrix organization (p=0.01) (**Figure 4D** and **4E, supplementary table 10**). Our findings suggest that PD-L1^+^ NK cells may traffic between lymph nodes and the peripheral blood to perform cell killing functions.

Next, we assessed whether this NK cell subset belongs to a specific maturation stage of NK cells (reviewed in ^27,28^). First, we repurposed the CITE-seq data to characterize the maturation stage of CD56^+^ PD-L1^+^ NK cells: we clustered NK cells based on their maturation stage including progenitor cells, stage 3, stage 4A, stage 4B, stage 5 and stage 6. Our results indicate that CD56+ PD-L1+ NK cells preferentially belong to NK maturation stage 5 and 6 (**Figure 4 B**). To confirm this observation, we developed a large flow cytometry Cytek Aurora panel composed of 26 markers (**Supplementary Table 2**) that was designed to capture all NK cell maturation stages (CD34^+^ hematopoietic progenitor (stage 1-2), stage 3, stage 4A, Stage 4B, Stage 5 and stage 6), and we included other relevant functional NK cell markers such as CD107a, CD103, NKp44, NKp80, NKG2A, NKG2C, KLRG1, CD62L, Granzyme B, Perforin, IFN-γ. We then tested this panel on eleven different participants: 5 younger (aged 18 to 31 years old) and 6 older (aged 61 to 83 years old) **Supplementary Table 1**). These data demonstrate that PD-L1^+^ cells belong to the NK maturation stage 5 and 6 (**Figure 4 B** and **Figure 5 A & B**) and are mature NK cells (**Figure 5 D**, PD-L1 and CD56). Investigation of relative expression of PD-L1 indicate a significant emergence of PD-L1^+^ expression among NK cells during maturation stages 3, 5 and 6 (p=0.02, **Figure 5 C, K & L**). Detailed manual gating of this cell subtype is shown in **Supplementary Figure 5**.

**Figure 5:**
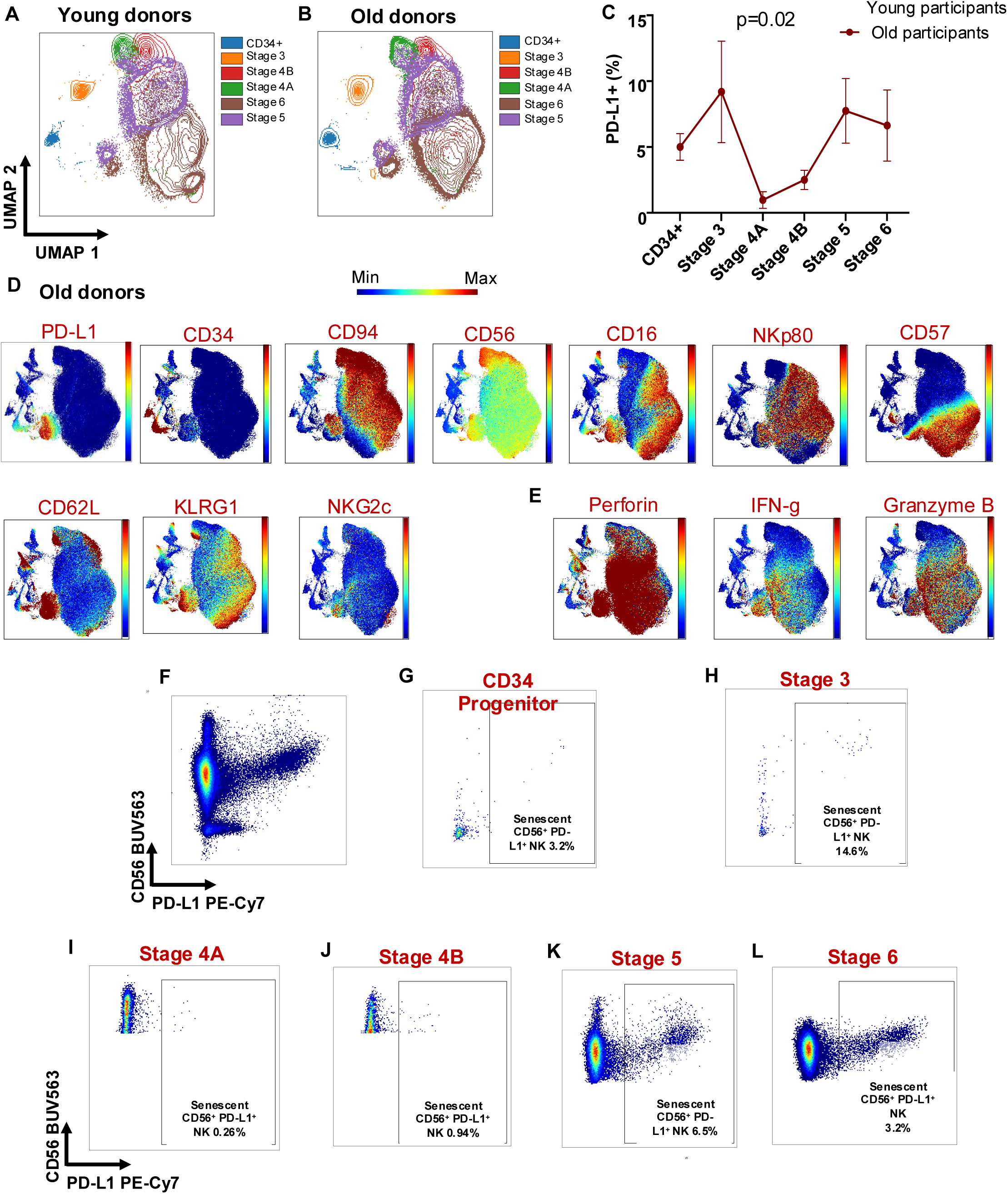
Phenotypic and functional characterization of the age associated CD56+ PD-L1+ NK cell subset. Cryopreserved human peripheral mononuclear cells (PBMCs) were thawed and stained with a large panel of cell surface markers and cytokines targeted to capture NK cell granularity. Samples were analyzed with a 27 colors Cytek Aurora panel (described on the methods). (**A**) Uniform Manifold Approximation Projection (UMAP) of NK cells representing the different maturation stages in young donors (n=5, <31 years old) and (**B**) old donors (n=5,>61 years old). (**C**) Using flow cytometry we measured changes in PD-L1 expression across the different NK cell maturation stages (CD34 progenitor cells, stage 3, stage 4A, stage 4B, stage 5 and 6). (**D**) UMAP projection of NK cells derived from 5 old donors, and showing expression of key NK cell differentiation markers (CD34, CD94, CD56, CD57, CD16, NKp80, CD62L, KLRG1, NKG2c) and (**E**) NK cell relevant functional markers such as Perforin, IFN-γ and Granzyme B. (**F** to **L**) Using NK cell panel Flow Cytometry data gated on live, CD45^+^ CD3^-^ lymphocytes, and on (**F**) CD56^+^ PD-L1^+^ NK cells. Next, CD56^+^ PD-L1^+^ NK cells were gated out of: (**G**) CD34 progenitor cells, (**H**) Stage 3 CD94^+^ CD117^+^, (**I**) CD16^-^ CD56^bright^ Stage 4A, **(J**) CD16^+^ CD56^dim^ Stage 4B, (**K**) CD57^-^ CD56^dim^ Stage 5, and (L) CD57^bright^ CD56^Dim^ Stage 6.

Further, we find that PD-L1^+^ NK cells co-express NKp80, a marker of functionally mature NK cells ^29^, CD62L and KLRG1 (markers of intermediate/mature and senescent NK cells ^12,30^), as well as CD94 and some NKG2C (**Figure 5 D**), which suggests previous exposure to CMV infection. Finally, we demonstrate that PD-L1^+^ NK cells exhibit cytotoxic features as well as perforin and granzyme B. They also display effector functions such as IFN-γ production (**Figure 5 D**). Overall, we describe an age-associated mature and cytotoxic NK cell subset distinguished by the co-expression of PD-L1 and CD56.

### “Immunological age” estimate and relevance to aging-related cognitive changes

In the field of senescence, multiple different “biological clocks” have been proposed with the aim of evaluating the difference between an individual’s chronological age and the state of that individual’s biological system given their genetic predispositions, environmental exposures and life experiences ^31–33^. Since immunosenescence strongly affects many important functions of the immune system and could influence a number of different health outcomes, we developed an estimate of a participant’s “immunological age” in an effort to initiate the translation of our results to the clinical sphere where cytometry data are readily available. Thus, we used elastic net regression (see **Method**) to select the most informative cell subtype frequency features with which to predict an individual’s chronological age (**Figure 6 A, Supplementary Table 5**). These cell subtypes are then used to estimate an individual’s immunological age. At the conclusion of this effort, our immunologic age explained 55% of the variance in chronological age, and we calculated a “delta age” for each participant (chronological age – immune age) (**Figure 6 B**).

**Figure 6:**
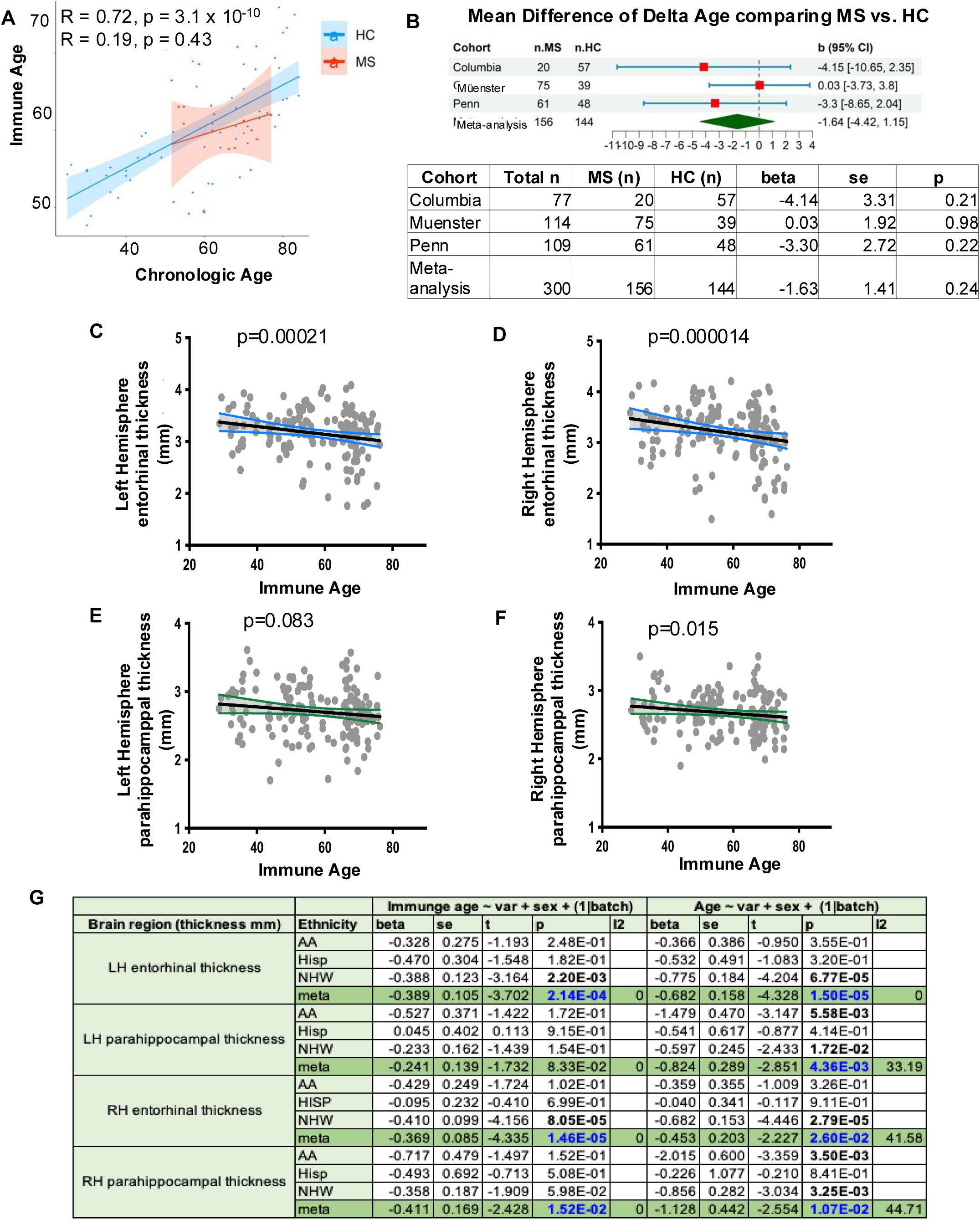
Immune age estimates and correlation with brain MRI measurements. Immune age is estimated using an elastic net regression model based on a leave one out cross validation model, considering the 14 cell subsets that are significantly associated with age. 55% of variance in chronological age, and (**A)** correlation with chronologic age. **(B**) Comparison of immune age estimates derived from healthy individuals with Multiple Sclerosis (MS) patients and summary table. (**C** to **F**) correlation of immune age estimated with MRI volumetric measures: (**C**) left entorhinal hemisphere (p=0.00021), (**D**) right entorhinal hemisphere (p=0.000014), (**E**) left parahippocampal hemisphere (p=0.08), and (**F**) right parahippocampal hemisphere (p=0.015). (**G**) Summary statistics Table correlating immune age estimate and brain volumetric measures.

To test the utility of our proposed “immunological age” estimate, we deployed it in participants with MS for whom we generated the same cytometric data as our healthy participants (demographics in **Supplementary Table 1**). Further, we repurposed two additional, existing datasets of MS and healthy participants from the University of Pennsylvania ^18^ and Muenster University ^19^. From these two independent cohorts we extracted 10 available cell populations associated with age (as defined in our analysis of RANN and Offspring participants), and we repurposed them to calculate an immunological age using the algorithm developed in our healthy participant data. We find that our algorithm captures a similar proportion of the variance in age in these other datasets, in healthy and MS participants (**Supplementary Figure 6**). We then calculated the delta age of each individual in all studies and compared the healthy and MS participants in each study. For the Columbia analysis, we only used the NHW healthy women participants since the MS participants were all NHW women. We then meta-analyzed the results of each of the three comparisons to provide the definitive evaluation of whether persons with MS display an alteration in their immune age. We find that participants with MS do not exhibit a significant difference in immune age (**Supplementary Figure 6**).

Next, since the role of the peripheral immune system in aging-related cognitive decline is an area of interest, we evaluated the relevance of our immunologic age measure to the central nervous system (CNS) by assessing whether it was associated with other phenotypic features available in the RANN participants, such as volumetric measures of brain structures derived from magnetic resonance imaging. We prioritized the entorhinal cortex and hippocampus, as they are the two regions implicated early in the course of Alzheimer’s disease, the most common cause of age-related cognitive decline. Interestingly, we find that left and right entorhinal thickness are associated with immunological age (**Figure 6 C & D**); further, left and right parahippocampi display suggestive evidence of association (**Figure 6 E & F**) in these participants who are cognitively non-impaired. Since the entorhinal cortex is the earliest brain region to be affected by Alzheimer’s disease and pathology then spreads to the hippocampus before going out to more distal neocortical regions, our results are intriguing. Further investigation of our estimate of immunological age will be needed to further disentangle the relationship of immunological age and regional brain atrophy that may be attributable to the earliest stages of AD.

### Trajectories of immunosenescence

To complement the analysis of immunologic age, we deployed the BEYOND approach that we recently proposed in the context of single nucleus RNA sequence data analysis to characterize trajectories of cognitive aging based on cross-sectional data ^34^. In short, we used our immunophenotypic data consisting of cell subtype frequencies collected by manual segmentation, and we projected each participant into a reduced dimensional space using the PHATE algorithm (**Figure 7 A**). Using a pseudo-time ordering approach (Palantir), we identified three distinct trajectories that emerge from the data; they start from the upper left corner of the graph and three different individuals define the end point of each trajectory (**Figure 7 B**). Each participant has a certain probability of belonging to each trajectory, and we refer to them initially as probability 1, probability 2 and probability 3 (one for each of the trajectories) (**Figure 7 B**). These probabilities are derived using the Palantir algorithm. The association of each of these probabilities to different traits is summarized in **Figure 7 C and Supplemental Tables 11** and **12**. Probability 1 displays a significant association with advancing immune age and we refer to it as “Immune Aging” trajectory (IA) **(Figure 7 B, E & G**). Probability 2 exhibits a stronger association with previous exposure to CMV virus (p=2.39 x 10^−10^) and advancing chronologic age (p=0.01) than probabilities 1 and 3 (**Figure 7 B, D, F and G**). Thus, we propose that probability 2 may capture a more general effect of aging, and we refer to it as the “Chronological Aging” (CA) trajectory. On the other hand, Probability 3 is quite different from the other two trajectories, and it is relatively depleted in terms of CMV exposure (p=1.64 x 10^−10^) (**Figure 7 B, F & G**) and significantly associated with younger age (both immune p=0.0002 and chronologic p=0.0001); hence, we refer to it as the “Low CMV” trajectory (**Figure 7 B, D, E and G**, **Supplementary Table 11**).

**Figure 7:**
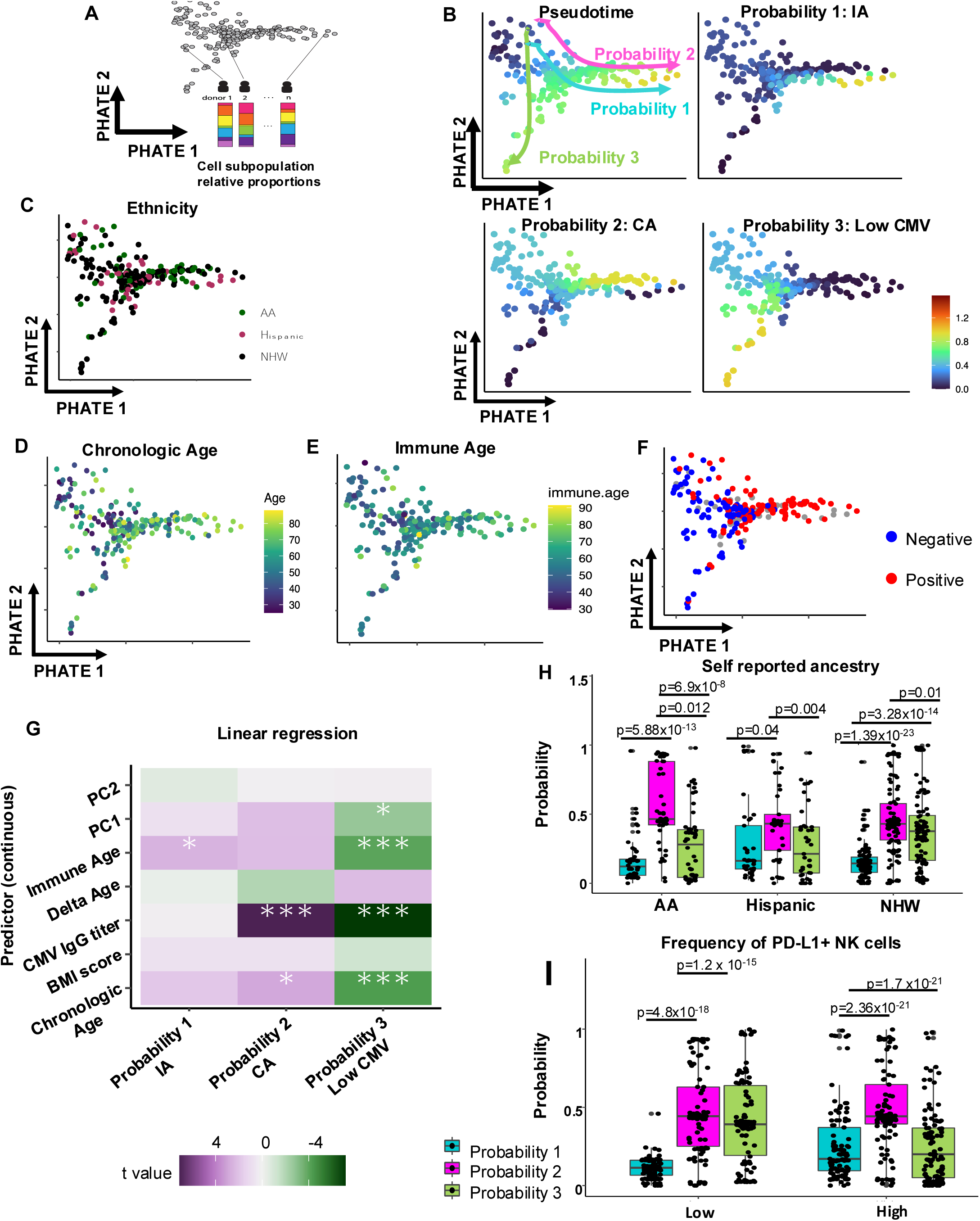
Trajectories of immunosenescence. **(A)** The structure of the participant’s immune age landscape across manifold of the lifespan. The age manifold captured by BEYOND algorithm for the 222 study participants. Demonstrating PHATE projection of each study participant (individual dot) based on similarity of the influence of environmental factors and ancestry background and represented as distinct aging trajectories (pseudotime). (**B**) Three distinct patterns (probability 1, 2 & 3) of immune aging trajectories across the diverse study participants manifold. Participants are colored based on the locally smoothed correlations with genetic and environmental factors (embedded within the landscape of aging trajectories), represented as pseudotime. PHATE Representation of distinct patterns of immune aging trajectories correlated with (**C**) Ethnicity, (**D**) chronologic age, (**E**) immune age, and (**F**) CMV serology. (**G**) Heatmap depict correlations between the three different probabilities with predictor variables, considering the different probabilities as the refence group. Based on significant associations, the probabilities are labeled as follow: probability 1: Immune Aging (IA), probability 2: Alternative Aging (CA), and probability 3: Low CMV. Predictor variables included: genetic ancestry (PC1and PC2), immune age, delta age, chronologic age, CMV IgG titer and BMI score. Significance is attributed based on t values, further statistics are provided within **supplementary Table 11**. (**H**) Representative plots highlighting the distribution of the three different probabilities across the self-reported African American (AA), Hispanic and non-Hispanic white (NHW) ancestry groups. Frequencies of CD56^+^ PD-L1^+^ NK cells were extracted based on the manual Flow Cytometry analysis. Participants were grouped as “low” if frequencies of CD56^+^ PD-L1^+^ NK cells were < 13.8%, or as “high” if frequencies of CD56^+^ PD-L1^+^ NK cells were > 13.8 %. (**I**) Representative plots highlighting the distribution of the three different probabilities within “low” vs. “high” CD56^+^ PD-L1^+^ NK cells participants.

Characterizing the role of the three different ethnicities of interest into our reduced dimensional space highlights that the African American group is preferably associated with the CA trajectory (**Figure 7 C** and **H**). Other traits such as hypertension **(Supplementary Figure 7 A)**, diabetes **(Supplementary Figure 7 B),** or BMI (**Supplementary Figure 7 C**) are not associated with these three trajectories (**Supplementary Table 11**). Next, we evaluated whether specific immune cell subsets that influence our estimate of immune-age impact these probabilities, and we demonstrate that PD-L1^+^ CD56^+^ NK cells are strongly associated with the CA trajectory (p=4.8 x 10^−18^, **Figure 7 I**), while the established age-associated immune cell subset of TEMRA CD8^+^ cells^3^ seems to strongly correlate with the IA trajectory (p=9.72×10^−27^) (**Supplementary Figure 7 E**). Overall, our findings suggest that exposure to CMV is a key factor that shapes trajectories of immune-aging over the lifespan, but not BMI nor hypertension, or diabetes. Genetic ancestry appears to have a modest contribution. Further investigation is necessary to study which other factors may trigger a faster immune aging or reverse it.

## Discussion

To date, existing data related to immune aging has been predominantly generated from populations of European decent utilizing cross-sectional study designs, and they employ mainly a categorical comparison of age groups (young *vs.* old) perhaps due to their smaller sample sizes ^3,35,36^. As a result, our understanding of immune aging trajectories and inter-individual variability across human populations remains limited. Understanding the physiology of ISC may enhance our ability to predict mortality risks and therapeutic outcomes in the elderly and in persons with neoplastic and chronic diseases.

Here, we report several biological and methodological insights. First, we found that known immune cell populations that are affected by aging, appear to largely behave in a similar manner among AA, Hispanic and NHW participants living in the same environment in Northern Manhattan: the I^2^ value in **Table 1** is mostly at or near 0, denoting minimal heterogeneity among the three groups partitioned based on self-reported race and ethnicity. There are some notable exceptions, including the interesting CD11c^+^CD20^+^ B cell population that is much more affected by age among AA participants than among other participants. This illustrates the fact that some of the effects of aging on the immune system are heterogeneous: they vary substantially even among different groups of individuals living in the same geographic area. Disentangling the role of genetics *vs.* the life experiences and environmental exposures which drive these ISC differences is challenging, and, importantly, we addressed these challenges by capturing both self-reported ancestry (a sociocultural construct) and genotype-derived estimates of genetic ancestry in most of our participants. The categories of self-reported ancestry are best modeled using a meta-analysis approach that respects the possible distinct characteristic of each group, and it allows for the calculation of a measure of heterogeneity among the three strata of data (I^2^). It is reassuring that the meta-analysis results are largely captured by the analysis using covariates that capture genetic ancestry, showing that both methods are reasonable. The genetic-based approach has the advantage that all participants can participate in the analysis, even those who report no self-reported ancestry or are of a different (Asian) ancestry. While self-reported ancestry and genetic ancestry are strongly correlated, they are not interchangeable and are used to ask different questions. The analysis reported in **Figure 3B** suggests that genetic ancestry has modest effects on the frequency of the cell populations in peripheral blood that we have measured; it is more likely that life experiences or environmental exposures would explain the heterogeneity that we are seeing amongst the three main populations that we studied in this report.

One example of such an environmental exposure that strongly influences ISC associations is infection with CMV. Consistent with prior studies ^37–39^, CMV has a strong effect on the proportion of certain cell subtypes, but it seems to affect the three groups (AA, H, NHW) in a similar fashion. Thus, it does not appear to be a driver of the inter-group heterogeneity that we observed. The analysis in which CMV is used as a covariate highlights the fact that the effects of aging and CMV are strongly intertwined, so that we cannot disentangle which of the two factors may cause the cellular changes. We will need longitudinal data to resolve this question: our cross-sectional data cannot resolve this issue of causality. Finally, the trajectory analysis highlights one trajectory that is quite distinct from the two dominant trajectories that are most associated with advancing age: the IA and CA trajectories. We label this third trajectory as the “Low CMV exposure” trajectory as the majority of the participants that define this trajectory do not display evidence of having been exposed to CMV. Since this third trajectory does not associate strongly with aging, it may represent an alternative path of immune function whose benefits and risks remains to be determined.

The CD56^+^PD-L1^+^ NK cell population that emerged from our analyses is intriguing in a number of ways. First, it is one of the cell populations that has the most heterogeneous effect, with an association with age that seems stronger in Hispanics than in other populations. It also seems to drive the terminus of the IA trajectory (**Supplementary Figure 7G**). Age-associated changes within the NK cell compartment have been reported previously, including changes within their phenotype and function as well as enhanced proportions of CD8^+^ CD56^+^ NK cells ^3,37,40,41^. Characterizing the diversity and nature of NK cell subsets is crucial for understanding several diseases: for instance, increased frequencies of CD56^bright^ cells have been documented in lupus erythematosus ^42^, in patients infected with hepatitis C ^43^, as well as in cancer patients ^44^. Additionally, the efficacy of daclizumab (anti-CD25) in MS is believed to stem from the expansion of immunoregulatory CD56^bright^ NK cells that migrate to the CNS and directly exert cytotoxic properties on autoreactive T cells ^45–47^ ^48^. Hence, to provide context to our data highlighting novel age related CD56^+^PD-L1^+^ NK cells, we elected to dive deeper into the characterization of this cell subset, examining gene expression, potential functions, and maturation stage. This NK cell subset exhibits a mature phenotype (stage 5/6), and PD-L1 expression appears to emerge during stage 5. It also displays other NK cell maturation markers such as CD62L (L-selectin) and the senescence marker KLRG1^30,49–51^. L-selectin is known to help trafficking of NK cells to lymph nodes and is associated with effector NK cell functions ^30,49^. While KLRG1 is known to regulate NK cell homeostasis and is known as an NK cell proliferation marker ^51^. Furthermore, we provide additional evidence that the CD56^+^PD-L1^+^ NK cell subset expresses granzyme B and perforin, suggesting a cytotoxic behavior, which is also supported by the presence of other effector functions such as IFN-_γ_ production. We also observed an enhanced expression of NKG2C associated with CD56^+^PD-L1^+^ NK cells, indicating earlier CMV infection ^52–54^. Future analysis will focus on characterizing the *in vitro* cytotoxic activities of the newly reported CD56^+^PD-L1^+^ NK cells.

We have developed an algorithm that leverages selected cell subpopulations to infer an individual’s immunologic age, and we have demonstrated that it is portable to other datasets of healthy individuals and persons with MS. Certain studies using epigenomic clocks have suggested evidence for accelerated aging in this chronic inflammatory disease ^55,56^, but the interpretation of these results remains unclear. Here, combining three different collections of persons with MS and controls, we do not see evidence for a significant alteration of ISC in MS (**Figure 6 B**). This analysis has some limitations, as we repurposed two independents datasets ^18,19,57^ that were generated for other purposes including data from untreated/treatment naïve persons with MS, and the three cohorts of persons with MS have different proportions of individuals with different levels of disability and disease stage, as evidence by differences in the number of participants with progressive MS. Thus, we cannot rule out a small difference in immunosenescence in the context of MS and/or that an effect may be limited to a subtype of persons with MS based on age, disease state or previous DMT exposure.

While the relation of immune age to an inflammatory disease such as MS requires further investigation, the relation of immune age to entorhinal cortex atrophy is more robust and intriguing. It joins other evidence ^58–60^ that there may be a relation between the peripheral immune system and aging-related brain atrophy and subsequent cognitive impairment. Prospective population-based studies suggest that regional brain atrophy occurs before overt cognitive decline and that the entorhinal cortex is among the first regions to display age-related atrophy ^61,62^, likely because it is the site where tau proteinopathy accumulates first and leads eventually to aging-related cognitive decline from the most common form of dementia, Alzheimer’s disease (AD). Our participants are not cognitively impaired, and thus entorhinal cortex atrophy and the more modest atrophy of the hippocampus (the next region to be affected by the stereotypical spread of tau proteinopathy) may be marks of the earliest stages of neurodegeneration linked to AD. Our measure of immune age is associated with this neurodegenerative process. Our cross-sectional data unfortunately do not yield insights into whether there is a causal relationship between these two traits (which requires longitudinal data), but it provides a robust foundation for future studies to explore this issue.

The inference of immune trajectories from cross-sectional data (**Figure 7**) is enabled by assuming that each participant represents one point along the different trajectories. Our modeling returns the most likely solution as being 3 trajectories. Once these trajectories are established, each individual has a probability of belonging to these trajectories, and we can relate phenotypes to these probabilities to see which traits may be related to one or more trajectory. Thus, we find that two of the trajectories are likely related to advancing age while a third trajectory is not strongly related to aging and appears to be defined by the absence of CMV infection. It is also negatively correlated with aging (both immune age and chronologic age), meaning that it seems to be more prevalent in younger individuals. Among the two aging trajectories, the CA trajectory seems to be enriched among AA participants, suggesting some heterogeneity in these trajectories amongst our 3 population groups. External data can be projected into these trajectories so that we can leverage our reference data to re-analyze external datasets.

There are several important limitations to our study. First, the absence of longitudinal data limits our understanding of individual changes in time compared to baseline data for a precise characterization of immune aging trajectories. While we excluded participants with inflammatory disease from the ISC analyses, it will be important to significantly expand our overall sample size to enhance our statistical power to conduct more advanced modeling and better define the role of common age-associated co-morbidities. Second, our estimate of immune age will need further validation leveraging independent, large cohorts prior to being tested in the clinical environment. Finally, we have not yet evaluated our ISC profile and trajectories in the context of cancer and its immunotherapeutic regimen; this area of investigation is likely to uncover strong influences of our immune age estimate and to establish the role of certain interesting cell populations like the CD56^+^PD-L1^+^ NK cells in rendering older individuals more susceptible to neoplasia.

In summary, our study explored immune-aging trajectories across diverse populations and explored environmental factors that may influence inter-individual immune heterogeneity. We uncovered an age-associated NK cell population co-expressing PD-L1, that exhibits a mature and cytotoxic phenotype. Additionally, we report that immune-age estimates are greatly influenced by previous CMV exposure and genetic ancestry. Furthermore, we identified a robust association between peripheral immune-aging and regional brain atrophy. Future work is warranted in a longitudinal study setting to expand our findings, and dissect whether CMV causes accelerated immune-aging, and in turn affects the brain functions and cognition.

## Acknowledgements

We thank all the study participants and Multiple Sclerosis (MS) patients who donated blood and participated in this study. We thank Drs. Yaakov Stern and Zonqi Xia who generously shared blood samples with us. We thank Drs. Bar-Or, Luiza Klotz and Heinz Wiendl for sharing data from MS patients. We thank Drs. Leah Zuroff, Koji Shinoda and Rui Li who generated the Penn MS dataset. We thank Drs. Hegewisch Solloa and Mace for the input regarding the NK cell maturation experimental design and data discussion. We thank Drs. Habeck and Stern for discussing the environmental factors and MRI data. We thank Dr. Gray and Farber for sharing their automated analysis pipelines. We thank Dr. Isnard for running the CMV ELISA experiments. H.T. received the merged Canadian Multiple Sclerosis Society (EGID 3855) and Fonds de recherche du Quebec en santé post-doctoral fellowship (292503), and the National Multiple Sclerosis Society Career Transition Award (TA-2305-41290). This work was supported by gift funding from the Ludwig Family Foundation (to P.D.L), NIH grants (R01 AG038465 to Y.S. and C.H., RF1 AG058067 and R01 AG054070 to J.J.M. and A.M.B.). The RANN and Offspring studies IRB numbers are: AAAI2752, and AAAR1120, respectively.

## Contributions

Project designed and funding obtained by P.L.D. Participant recruitment Y.S., Z.X, and P.L.D. Assay development and miniaturization H.T. Blood sample processing J.M. Data acquisition H.T., A.K. and C.H. Data analysis H. T., A.L., T.L., M.F., N.C.L., C.H., and P.L.D. L.Z. and A.B.O. informed design of PBMC and T cell flow cytometry panels and shared Multiple Sclerosis data. NK cell panel design and data discussion H.T., E.H.S and E.M. CMV ELISA S.I. and J.P.R. A.B.O., L.Z., H.W. and L.K. shared Multiple Sclerosis data. T.L., M.F. and V.M. helped implementation of automated flow cytometry analysis. A.B.O., L.K., Y.S., E.M. discussed findings. Manuscript writing H.T. and P.L.D. Final manuscript review & editing: all co-authors. All authors read and approved the final version of the manuscript.

## Material and methods

### Participants and ethics

A total of 222 adults without chronic conditions were recruited part of the immunosenescence (ISC) cohort, including n=205 participants from the RANN (Reference Ability Neural Network, IRB number: AAAI2752) study, and n=20 participants from the Offspring (Offspring of Washington Heights-Hamilton Heights Inwood Columbia Aging Project, IRB number: AAAR1120) study. All individuals were between 25 and 88 years old. Demographics details can be found in **Supplementary Table 1**. Pacific islander, Asians, biracial and other participants were not included in the self-reported ethnicity analysis due to the small numbers, reducing the final cohort to 187 participants. Three participants did not meet the quality control criteria and were left out of further analysis.

All participants provided an informed written consent form using protocols approved by Columbia University’s institutional ethics review boards. RANN Participants underwent brain imaging using magnetic resonance imaging (MRI) 3 Tesla machine. Older RANN individuals with dementia and mild cognitive impairment (dementia rating scale of <130); and subjective functional impairment (BFAS > 1) were excluded from this study. We also excluded participants with: heart diseases including congestive heart failure, stroke, CNS infections and tumors, epilepsy, Multiple Sclerosis, Parkinson’s disease, Huntington’s disease, head injury (loss of consciousness over 5 minutes, seizure, intellectual disability, normal pressure hydrocephalus, essential/familial tremor, listening disability such as dyslexia, ADHD or ADD, color blindness and uncorrectable vision, uncorrectable hearing and implant as well as participants receiving a CNS targeting drug, major depression, bipolar or psychosis episodes (past 5 years). We further excluded participants with down syndrome, AIDS, uncontrolled thyroid, or other endocrine related diseased, pregnant, and lactating participants, or participants with untreated neurosyphilis, with alcohol and drug addiction within the last 12 months. Participants with a history of cancer within the last five years, renal insufficiency, active hepatic disease, non-skin neoplastic disease/melanoma and insulin dependent diabetes. Some participants were under cholesterol, hypertension and diabetes disease controlling therapies ^21^, which reflects the demographics of a normal aging population. RANN participants were required to be native English speakers, strongly right-handed and have at least a fourth grade-reading level.

Offspring participants are the children of WHICAP study participants and residents of northern Manhattan, New York at the time of the study ref. Participants did not have a diagnosis of dementia, underwent a psychological evaluation, self-reported their race and ethnicity. At the time of recruitment, history of diabetes, heart disease, stroke and hypertension are ascertained by self-report. All Offspring participants have signed a consent approved by the institutional Review Board at Columbia University^63^. A total of 159 untreated MS patients were recruited part of the study, including 20 patients recruited at Pittsburg university aged between 51 and 77 years old, one patient recruited at Columbia University (54 years old), 78 untreated MS (38 RRMS and 40 PPMS) with 44 age and sex matching controls were recruited and phenotypically characterized at the University of Pennsylvania ^18,57^, and 61 untreated MS (37 RRMS and 24 PPMS) with 50 controls were recruited and phenotypically characterized at Muenster University^19^.

### Study design

The overall aim of this study is to shed light on peripheral immunosenescence trajectories across a heterogeneous population, leveraging advanced Flow Cytometry analysis approaches to extract cellular features significantly associated with age. Using the age associated cellular features, we developed an immunological age estimate model. In addition, we aimed to correlate age associated cell subsets with different brain region thicknesses. Peripheral blood mononuclear cells (PBMC) were thawed in batches of 8 participants, and over a period of 8 months. Immune cell subset phenotyping analysis was conducted in a batch-batch manner. Experimental batches were designed in a balanced manner (age, sex, ethnicity), including 8 participants each (female: male 4:4, young: middle aged: old 2:4:2 and a mix of all ethnicities). A participant was used as an internal control sample every other month across all batches.

RANN participants first underwent cognitive tests outside of the scanner using the American version of the National Adult reading test (AmNART) ^64^ to estimate IQ. Moreover, four cognitive domains including reasoning, vocabulary, memory, and speed of processing were created. A total score of each cognitive domain was derived by computing the average z-score of the cognitive test grouped within the same factor. Next, brain imaging MRI brain volumetric measures included structural measures, reginal brain volumes and cortical thickness were obtained from submillimeter resolution T1-weighted MRI data. Results were processed using FreeSurfer’s 5.1 pipeline ^65^, and further implemented to detect small changes in the gray matter. This is an overall robust processing pipeline but could be subject to minor variation in the optimization process ^66^. Despite the automated process of the estimation, we manually validated the accuracy of the spatial registration, white and gray matter segmentation were manually verified, following the analytic procedure as outlined Fjell et al., ^67^.

### Sample processing

Peripheral venous blood was collected (40mL) from a total of n=222 participants part of the ISC cohort and n=20 persons with MS. PBMC were enriched and cryopreserved for up to 3 years using strict standard procedures (SOPs) developed and validated by the Center of Translational and Computational Neuroimmunology at Columbia University Medical Center, and as previously described in ^68^ using density centrifugation and Lymphoprep (STEM Cell Technologies, Canada). Samples were retrieved from the liquid nitrogen tank and transported to the water bath in dry ice. PBMC vials were thawed in the 37 °C water bath for 1-2 minutes and resuspended in warm X-VIVO 10 serum-free cell-media (Lonza). Samples were washed by centrifugation for 12 minutes at 1200 rpm at room temperature (RT). Cell viability was assessed using the automated cell counters at Nexcelom (Nexcelom, Bioscience) with acridine orange and propidium (AOPI) viability dye (Nexcelom, Bioscience). Sample viability was confirmed using the fixable viability dye as per manufacturer recommendation (LIVE/DEAD Aqua Cell stain kit, Thermofischer Scientific) and measured using flow cytometry (BD LSR II, BD Biosciences). The median cell count of thawed PBMCs is (4 million cells per participant), and the majority of samples had viabilities ≥ 90%, while samples with a viability < 70% (n=3) were excluded from further analysis. Demographics, cell counts, and cell viability of each participant are described in **Supplementary Table 1**.

### Immunophenotyping of cryopreserved PBMCs

Using classic multiparametric flow cytometry to interrogate PBMCs part of the ISC cohort, we deployed two panels designed to phenotypically characterize PBMCs and T cells. Panel design was in part informed by ^18^ and validated using rigorous standard operating procedures (SOPs) adapted from Dr. Bar-Or’s team ^18^. We emphasized subsets with known or hypothesized relevance to natural senescence processes, while adopting computational analysis approaches to capture novel immunosenescent associated immune subsets. To further phenotypically and functionally characterize the novel NK cell subsets we designed a panel composed of 28 cell surface and intracellular markers designed to capture NK cells across the different maturation stages, and with effector and cytotoxic functions using Cytek Aurora machine (Cytek Bioscience).

We assessed a total of 8 x10^5^ – 1 million PBMCs for flow cytometry experiments and used 96 v shape plates for further cell staining. Prior cell surface staining, PBMCs were stained with the LIVE/DEAD Aqua viability dye for 30 minutes protected from the light at room temperature (RT) and washed twice using cold phosphate-buffered saline (PBS) and resuspended in cold PBS. For the cell surface marker staining we used strict SOPs developed by ^69^, and used manufacturer recommendations (BD Biosciences). Briefly, we performed cell surface staining using flow cytometry (FACS) buffer (PBS containing 2% FCS + 0.1 mM EDTA). Cells were suspended in buffer containing a pre-mix combination of fluorochrome-conjugated antibodies and incubated on ice for 20 minutes (FACS buffer). For immunophenotyping using the LSR II Fortessa we used a 15-color panel designed to capture PBMCs and a 12-color panel specific for T cells. Aurora was utilized to further characterize NK cell subsets. All cell surface markers used in this study are described in **Supplementary Table 2**. Stained cells were washed twice with cold FACS-buffer to remove unbound antibodies and fixed in 2% paraformaldehyde (PFA) for further flow cytometry analysis. Unstained sample controls for each donor were processed alongside experimental samples.

All flow cytometry sample acquisition was performed by the same operator who was blinded to the sample source cohort and followed strict standardized protocols for acquisition. Sample acquisition was performed using LSR II flow cytometer (BD Bioscience), and Cytek Aurora (Cytek Biosciences).

### Flow Cytometry analysis

To investigate novel immune cell subsets associated with age, we deployed a high-dimensional R based flow cytometry data analysis approach as informed by ^70^, and compared to the classic bi-dimensional FlowJo analysis for further approach validation.

### Quality control

For high-dimensional flow cytometry analysis, we used the classic manual bi-dimensional gating strategy using FlowJo 10.7 (Becton Dickinson BD, NJ) to pre-process the flow cytometry data. First, we manually adjusted compensations, then gated out cellular debris, doublets, granulocytes and dead cells. Representative examples of staining, quality control gating strategy and gating of major lineage immune cells such as T-cell subsets, B-cell subsets, myeloid cell subsets and NK cell subsets are shown in **Supplementary Figure 4**. Cleaned data was exported as .fcs files of compensated live cells and further imported into Seurat R package (version 4.1.1) to assess their quality and run-in depth analysis using the computational pipeline. Cells over the 90^th^ decile of amine-reactive dye fluorescence were labeled as dying. Principal Component Analysis (PCA) on various summary statistics were performed to identify outlier samples with higher proportion of dead cells and a lower number of cells, which were further excluded from downstream analyses. To further assess the quality of the dataset, flow rate, signal acquisition, and dynamic range were assessed using FlowAI to detect and remove anomalous cells ^71^.

### Transformation and clustering

To better capture the wide range of fluorescence, we assessed the ability of two common transformations, logicle and asunh, to merge cells from the ISC cohort (n=247) using PCA and Uniform Manifold Approximation and Projection (UMAP). As a reference, we use the untransformed fluorescence values. Lgicle performed better; therefore, we used it for downstream analyses and visualizations.

We clustered the filtered and logicle-transformed fluorescence matrices were clustered with PhenoGraph using Euclidean distance to build a network with cells as nodes and Jaccard similarity as edge weights ^26^. Louvain community detection was then run iteratively until modularity did not increase in 20 iterations, a time limit of 2000s was reached, or 100 iterations had passed ^26^. To evaluate the robustness of the clustering solution, we trained a random forest with 500 trees to classify cells based on PhenoGraph’s communities using the prediction error and visual inspection of the confusion matrix to assess performance. Cells were plotted on the two-dimensional UMAP, and colored based on the clustering solution. To keep only high confident clusters, we excluded clusters with more than 10% of cells coming from one sample to keep only high-confidence clusters.

### Associations

We used participant level and cell level associations to detect cell clusters associated with age. First, we calculated cell proportions per participant and per cluster. Next, we applied logit transformation to liberate cell proportions from the 0-1 scale. To account for 8 participants per batch, we used random intercept for batch and fixed effects for age, sex and the first three principal components of genetic ancestry (PC1, PC2 and PC3).

an: a sample-level association and cell-level association. First, we calculated the cell proportions per subject per cluster and then applied log transformation to liberate proportions from the 0-100 scale. Then, transformed proportions for each cluster were corrected for the batch effect using the following mixed effect model, thus accounting for the fact that multiple samples were processed in the same batch:

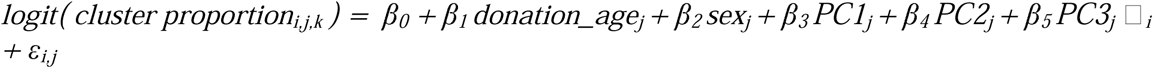

Where *i* indexes batches, *j* subjects, *k indexes clusters,* β*_0_* is the intercept estimate, □*_i_* accounts for multiple samples belonging to the same batch, and ε*_i,j_* is the error term. We also ran two additional models in individuals from the RANN cohort to model cytomegalovirus seropositivity and titers. We used the variance inflation factor to ensure multicollinearity was not present in final models.

For the cell-level association, we used the formulation by ^72^, where cells belonging to each cluster were labeled as 1 or 0 otherwise, and a mixed effect logistic regression model was fit to account for subject and batch variability as random effects. Age, sex, the first three principal components of genetic ancestry were modeled as fixed effects. We used the R package lme4 and the bobyqa optimizer ^73^to fit the following model:

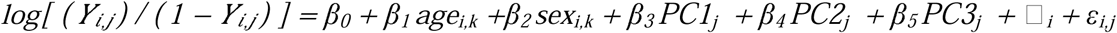

where *Y_i,j_* is the odds of cell *i* belonging to cluster *j*, β*_0_* is the intercept for each cluster *j*, □*_b_* is the random effect for each *b*th batch, and □*_k_* is the random effect for each *k*th subject.

Only terms below a false discovery rate of 5% for both modeling strategies were reported. For regression plots, we used the R package ggeffects to compute marginal means for the response variable while holding the other covariates at either the control level or the average value over the levels of factors ^74^.

### Estimation of genetic ancestry

Genotyping was performed for participants of the Reference Ability Neural Network (RANN) cohort (two batches: *N* = 429 and *N* = 142) by using single nucleotide polymorphism (SNP) array. Genotype data were separately compared with the TOPMed reference panel ^75^ (and cleaned by using the imputation preparation tool (version 4.3.0; https://www.well.ox.ac.uk/∼wrayner/tools/). Cleaned genotype data were uploaded to the TOPMed imputation server (https://imputation.biodatacatalyst.nhlbi.nih.gov/) and imputed with Minimac4 and the TOPMed reference panel (version R2). SNPs with imputation quality *r*^2^ < 0.3 were filtered out. Imputed genotype data were merged using PLINK version 1.90. SNPs were excluded if they had minor allele frequency < 5%, *p*-value of the Hardy-Weinberg equilibrium < 1 × 10^-5^, or non-zero missing rate. Linkage disequilibrium-based SNP pruning was performed with window size 250 kb, step size 10 kb, and *R*^2^ < 0.05, which left 48,695 SNPs. Genetic ancestry was estimated by using ADMIXTURE version 1.3.0 ^76^. The numbers from 1 through 6 were specified as the number of ancestral populations *K*. Comparison of estimated genetic ancestry with self-declared race and ethnicity showed that the results with *K* = 3 is the most interpretable one. Five-fold cross-validation also showed that *K* = 3 is sufficiently robust. Therefore, we employed the genetic ancestry estimates with *K* = 3 for covariates of association testing.

### Assessment of CMV IgG titers

Serum samples were collected during the same blood draw as the visit collecting PBMC for the cytometric characterization from each RANN cohort participant (n=208). Anti-CMV IgG levels were measured using immunoassay (ELISA) test kits (GenWay Biotech, San Diego, CA, USA). Serum samples with anti-CMV IgG levels greater than 0.25 IU/mL were considered CMV seropositive, and those with anti-CMV IgG levels lower than 0.25 IU/mL were considered CMV seronegative. For each assay we used a donor as a positive control. See **Supplementary Table** 4 for CMV IgG titers.

## Statistical analyses

### Association between chronological age and manually segmented cell subsets among individuals of different self-reported ancestries

To detect the impact of chronological age on the frequency of immune cell subsets, we first transformed the frequencies of immune cell subsets curated using manual FlowJo analysis to follow a normal distribution using Tukey’s Ladder of Powers in R package *transformTukey* and orderNorm Transformation in R package *bestNormalize*. Since we considered three self-reported ethnicity total (n=189), African-American non-hispanic, Hispanic, and non-Hispanic White, we tested the association of the normalized immune cell subsets with age by fitting a linear mixed-effects regression model adjusted for sex as fixed effect and batch as a random intercept in each of the three groups separately using R package *lme4* and combined the results using a random-effects meta-analysis using R package *metafor*. For comparison, we considered a “naïve model” in which we analyzed all three groups of participants together using a single linear mixed-effects model adjusted for age as fixed effect and batch as a random intercept. We further adjusted for the first three principal components (PCs) in the model to test the association of the normalized immune cell subsets with age using a linear mixed-effects model adjusted for sex and first three PCs as fixed effects and batch as a random intercept. For the RANN participants, we tested the association of the normalized immune cell subsets with age by fitting a linear-mixed effects model adjusted for sex, first three PCs, and CMV status (CMV seropositive or seronegative) as fixed effects and batch as a random intercept.

We further investigated whether the association between chronological age and immune cell subsets vary by sex, ethnic group, or MS status. To account for differences between experimental batch effects, a linear mixed-effects model was fitted on the normalized immune cell subsets adjusted for batch effect as a random intercept. We used the residuals from the model which represent a quantitative trait capturing variability in normalized immune cell subsets not captured by known technical factor. We tested the interaction effect between age and a) sex, b) ethnic group, and c) MS status on the residuals by fitting a linear regression model on the residuals and generated plots within **Figure 2** and statistics summarized in **Supplementary Table 1, 5 & 6**.

We investigated whether the effect of the CMV on normalized immune cell subsets would be mediated by chronological age. We also investigated whether the effect of chronological age on the normalized immune cell subsets would be mediated by CMV. We used a causal mediation analysis aimed at identifying whether the chronological age resulted from CMV or the reverse. In the mediation analysis, the mediated effect is the effect of the exposure on the outcome that occurs through a mediator variable. The direct effect is the effect of the exposure on the outcome after accounting for the mediator. The total effect is the sum of the direct and indirect effects of the exposure on the outcome.

### CITE-Seq

To assess the single cell transcriptome and proteome profile of the RANN cohort, we used the Total-Seq C Human Universal Cocktail consisting of 130 antibodies and 7 isotype controls (Biolegend, USA). Cells from 4 donors were pooled (150K Cells per donor), and the pooled cells were loaded into the 10X Genomics Chromium platform after cell staining. Briefly, after pooling, cells were centrifuged at 300g for 10 minutes at 4 °C, supernatant was discarded, and pooled cells were resuspended in Fc block staining buffer (Biolegend) in 5mL polystyrene flow cytometry tubes (Falcon, ThermoFisher, USA) and incubated for 10 minutes at 4 °C. The antibody pool (Total-Seq C Human Universal Cocktail, Biolegend) is added to the pooled sample in FcBlock/staining buffer and incubated for 45 minutes at 4 °C. 2mL of staining buffer are added to the pooled cells and samples are centrifuged at 300g for 5 minutes at 4 °C. The supernatant is carefully removed, and pooled cells are washed again. Finally, pooled cells are resuspended in x-vivo 15 (Lonza), filtered, counted and resuspended at a density of 2 500 cell/μL) and all cells are loaded into the 10x Genomics instrument.

### 5’ GEX, CITEseq and T cell receptor and B cell receptor sequencing library construction

scRNA-seq was performed using the 10x Genomics Chromium Single Cell 5’ Reagent Kits (v2 Chemistry Dual Index) with Feature Barcoding technology for Cell Surface Protein and Immune Receptor Mapping for simultaneous measurement of cell surface proteins along with the gene expression and immune repertoire information from the same single cell. Briefly, an estimated 20,000 CITE-seq cocktail antibody-labeled cells (pooled from 4 donors, as described above) were loaded on the 10X genomics chromium controller single-cell instrument. Reverse transcription reagents, barcoded gel beads, and partitioning oil were mixed with the cells for generating single-cell gel beads in emulsions (GEM). After the reverse transcription reaction, GEMs were broken, 10x Barcoded first-strand cDNA from polyadenylated mRNA and DNA from cell surface protein specificity Feature Barcode were amplified. Amplified full-length cDNA from poly-adenylated mRNA was used to enrich full-length V(D)J segments (10x Barcoded) via PCR amplification with primers specific to either the TCR or BCR constant regions. Amplified full-length cDNA from poly-adenylated mRNA was used to generate 5 Gene Expression library. Amplified DNA from the cell surface protein Feature Barcodes derived from the antibody was used to construct the Cell Surface Protein library. Independent constructed libraries were then analyzed and quantified using TapeStation D5000 screening tapes (Agilent) and Qubit HS DNA quantification kit (Thermo Fisher Scientific).

#### Sequencing

Independent sequencing libraries were pooled and sequenced on the Illumina NovaSeq 6000 with S4 flow cell according to the standard manufacturer’s protocol (Illumina, San Diego, CA) using paired-end, dual-index sequencing with 28 cycles for read 1, 10 cycles for the i7 index, 10 cycles for the i5 index, and 90 cycles for read 2.

#### Data preprocessing

FASTQ files were preprocessed with the CellRanger software (v6.0.0, 10x Genomics). To infer original participants for cells, we took advantage of single nucleotide polymorphisms (SNPs) in the RNA-seq reads. Cells in the filtered Unique Molecular Identifier (UMI) count matrix were grouped into four clusters, each of which represents one participant, based on RNA genotypes using the freemuxlet software (https://github.com/statgen/popscle). Assignment of clusters to participants were performed by comparing RNA genotypes in freemuxlet VCF file with DNA genotypes using Picard CrosscheckFingerprints.

#### Analysis of differentially expressed genes (DEG)

To assess whether PD-L1 is expressed by the age-associated CD56^+^ NK cell subset and not all NK cell clusters, we used the CITE-Seq data to run differential protein abundance analysis. We computed pseudobulk UMI counts of antibody-derived tag libraries for human peripheral blood cells. We compared differentially expressed genes and proteins in PD-L1^+^ CD56^+^ NK cells (cluster 13) to NK cells that express very low to no PD-L1 levels (Cluster 2 and 22). After the trimmed mean of M normalization, we applied the limma/voom test using seven isotype control antibodies included in the CITE-seq panel as covariates with FDR corrected threshold for significance at 0.05. The results of this analysis were visualized using volcano plots and showing FDR values <0.05 for significant differentially expressed genes (**Figure 4 C** and **Supplementary Figure 5 H**).

### Pathway analysis

To investigate gene pathways involved in the regulation of PD-L1^+^ NK cells (cluster 13) compared to NK cells that express very low to no PD-L1 levels (Cluster 2 and 22), we performed a gene set enrichment analysis using the reactome database ^77^, using the R package “clusterProfiler” with an FDR corrected threshold for significance at 0.05.

The results of this analysis were visualized using bar plots showing normalized enrichment scores and enriched gene pathways in cluster 13 compared to cluster 22 (**Figure 4 D**) or compared to cluster 2 (**Figure 4 E**).

### Deriving an estimate of an individual’s “immunological age”

To estimate a participant’s immunological age, we used an elastic net regression model to regress immune cell subsets on chronologic age in non-Hispanic White controls (n=124) adjusted for sex and batch to select the most informative features in predicting an individual’s immunological age. The model trained in non-Hispanic White controls was then used to estimate immunological age in Hispanic controls and African American non-hispanic controls, and non-Hispanic white untreated persons with MS. To test our model in a disease population, we used the 18-age associated immune cell subsets from external datasets derived from untreated and treatment naïve MS patients. These datasets included data collected at The University of Pennsylvania (Penn) n=109 and Muenster University n=114. Estimated immune sage were obtained after training the model on established control immune age estimates (ISC Columbia university participants). Established immune age estimates trained using Columbia university’s NHW controls was used to estimate the immunological age within the Penn controls (n=48) and MS patients (n=61) groups, as well as with the Muenster University controls (n=39), and MS patients (n=75) groups.

### Using Potential of heat-diffuser for affinity-based transition embedding (PHATE)

To understand the immunological age trajectories across different ethnicities, PHATE was performed in R using the phateR package ^78^ to align participants in a tew-dimensional space based on their immune-cell composition. The input to PHATE is a matrix of cell-type frequencies (z-scored across participants for each cell type). The input cell-type frequencies were estimated based on manual gating of markers for 25 cell-types across 219 participants. The parameters used in generating the PHATE map include principal components (n pca)=13, k-nearest neighbor (knn) = 5, kernel decay rate (decay) = 100, informational distance (gamma) = 0.5, diffusion power (t) = 24, distance metric for building knn graph (knn.dist.method) = Euclidean and distance metric for multidimensional scaling (mds.dist.method) = cosine. To visualize the patterns of phenotypic (e.g. immune age, CMV serology etc.) or demographical (e.g. age and ethnicity) variables across individuals in the same PHATE projection map. Each colored dot represents a study participant colored based on the variable inputted into the PHATE model (**Figure 7** and **Supplementary Figure 7**).

### Pseudotime Analysis using Palantir

Palantir was performed in Python using the Palantir package ^79^ on the RANN cohort. Conceptually, Palantir models immune cellular aging as so stochastic process, where a participant may traverse from one point to the next along the trajectories of immune aging with certain probability. Based on this framework, Palantir also determines the terminal states of the trajectories and assigns each participant a probability to reach a given terminal states. Computationally, Palantir first calculates a diffusion map based on the z-scored cell-type frequencies, using k-nearest neighbor (knn) = 9. The first seven diffusion components (n_components) were then input to the Palantir algorithm (run_palantir) to calculate pseudotime with knn = 9. The youngest individual was set as the starting point. To determine variables that drive a certain trajectory, linear regression was performed to link the probability of each trajectory to chronologic age, immune age estimates, ethnicities, CMV serology, hypertension and diabetes data were considered as model variables.

### MRI measurements

We collected structural brain measures from (n=167) RANN study participants, using Magnetic Resonance Imaging (MRI), using 3 Tesla machine. This approach enabled us to collect brain volume and cortical thickness - Submillimeter resolution T1-weighted MRI data, further processed with FreeSurfer’s pipeline (http://surfer.nmr.mgh.harvard.edu/). This approach is implemented to detect small to subtle changes in gray matter over time. The processing pipeline deployed here is robust to initialization points, which may lead small variations in the results of the optimization processes ^66^. Although the estimation procedure is automated, the accuracy of the spatial registration and the white matter and gray matter segmentations is manually verified following the analytic procedures described by Fjell et al ^67^.

### Immune age association with brain volumetric measurements

Brain volume thickness measures were associated with either immune-age estimates or chronological age of each RANN participant with MRI volumetric measure data (n=185). Briefly, immune age estimates or chronologic age were correlated with each brain region volumetric measure by fitting a linear mixed-effects regression model adjusted for sex as a fixed effect and batch as a random intercept in each of the three groups separately using R package *lme4* and combined the results (meta-analysis) using a random-effects meta-analysis using R package *metafor*.

## Figure legends

**Supplementary Figure 1:**
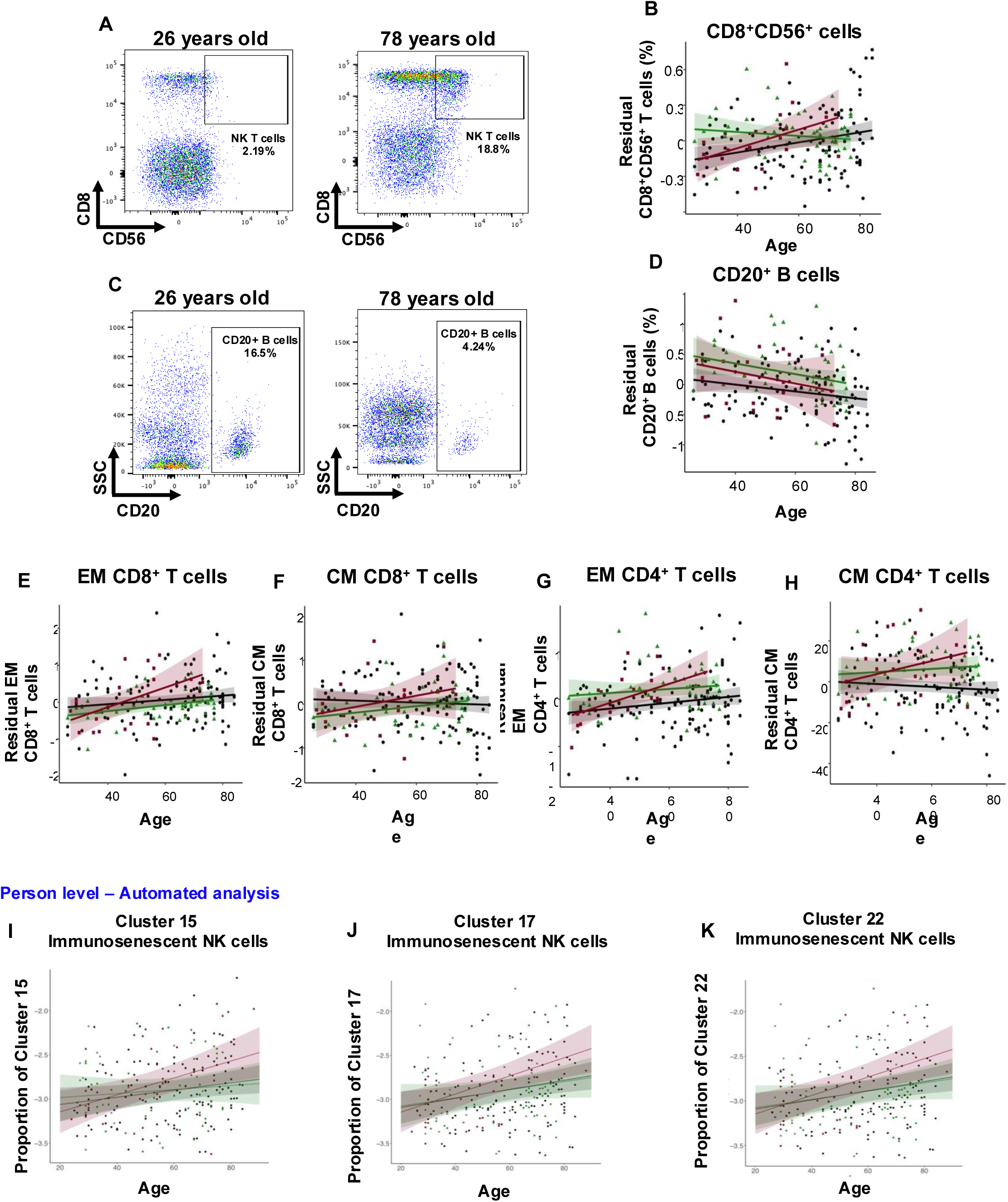
Comprehensive association between chronological age and immune cell subsets among individuals of different self-reported ancestries. Cryopreserved human peripheral mononuclear cells (PBMC) were thawed and stained with cell surface markers. Cell frequencies (%) are extracted using serial manual gating using FlowJo. Gating strategy is detailed in **supplementary Figure 4**. Representative Flow Cytometry plots of a 29 and a 69-year-old individual highlighting (**A**) CD56 co-expressing CD8 cells, and (**C**) CD20^+^ B cell subset. **(B**) Plot demonstrating the visual representation of linear regression model representing a positive correlation between CD56^+^ CD8^+^ cell frequencies with advancing chronologic age, driven by Hispanics (burgundy). **(D**) Plot demonstrating the visual representation of linear regression model representing a negative correlation between CD20^+^ B cell frequencies with advancing chronologic age. Plot demonstrating the visual representation of linear regression model representing a negative correlation between (**E**) EM CD8 T cells, (**F**) CM CD8, (**G**) EM CD8 and (**H**) CM CD4 T cells with advancing age. Detailed statistics are shown in **Table 1**. (**I** to **K**) .fcs Flow Cytometry files are analyzed in an automated approach using PhenoGraph and R package, and further linear regression models are deployed to assess correlations between immune cell clusters with chronologic age. Three distinct CD56^+^ PD-L1^+^ NK cell clusters are detected using the automated analysis (**I**) cluster 15, (**J**) cluster 17 and (**K**) Cluster 22, data projected at the cell level. Summary statistics are represented in **Supplementary Table 7.**

**Supplementary Figure 2:**
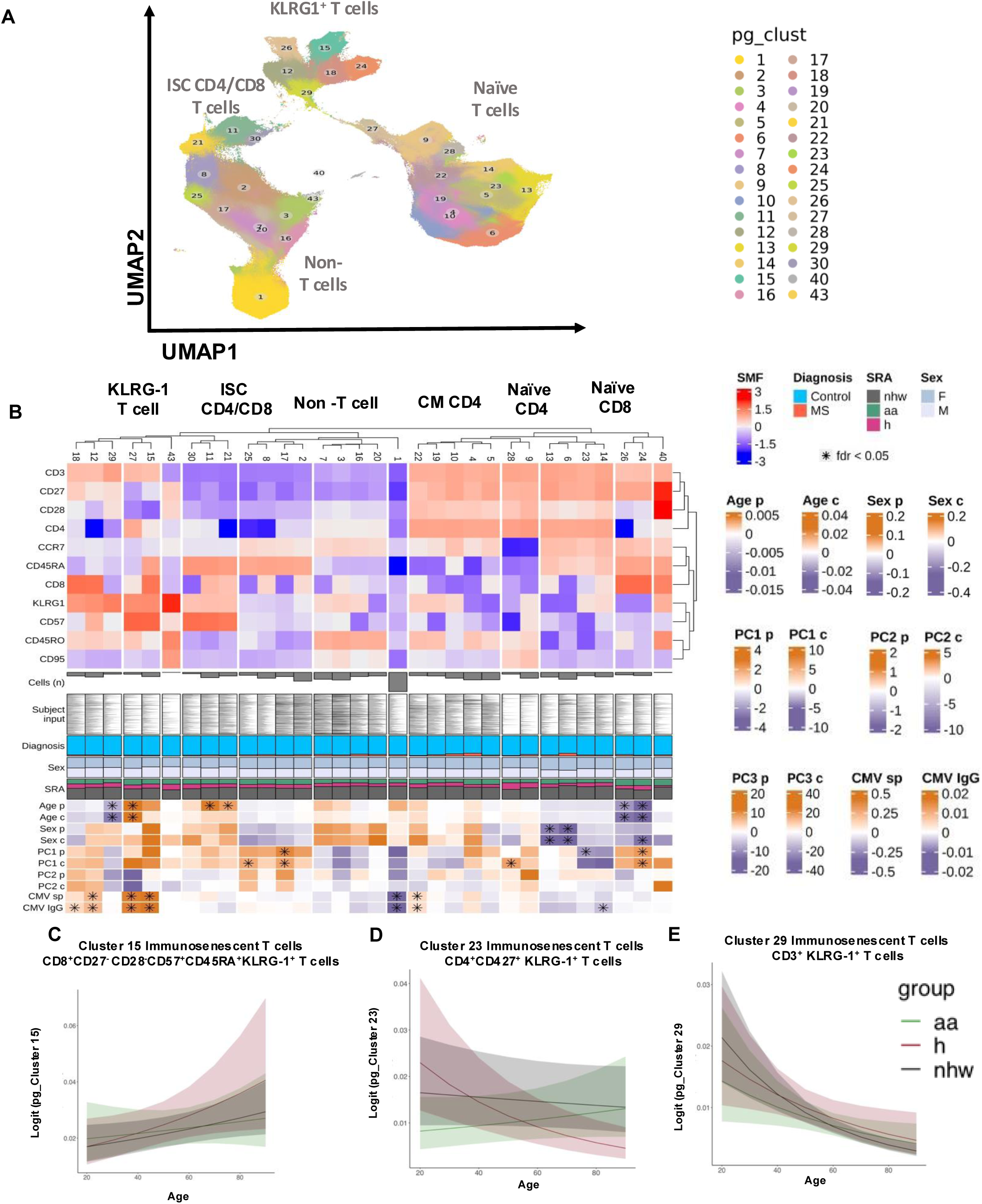
Automated cell clustering reveals novel cell clusters associated with chronological age among individuals of different self-reported ancestries. (**A**) Uniform manifold approximation and projection (UMPA) visualization plot of 32 clusters within the T cell panel based on the cell surface panel markers (CD3, CD4, CD8, CD27, CD28, CD45RA, CD45RO, CD57, CCR7 & KLRG1). (**B**) Heatmap depict fluorescence of each cell surface marker that help define the different immune cell subsets within 247 individuals (African American (AA), Hispanics and non-Hispanic whites (NHW)), and 21 untreated RRMS patients. Fluorescence ranges from −4 (blue) to +4 (red). Cell surface markers are used are presented across the top y-axis. Cell cluster numbers representing distinct cell subsets are ordered on top of the heatmap. The number of cells (n cell) within each cell cluster is expressed in the x-axis. The proportion (prop) of each cell subset across each cell cluster is represented in the x-axis. The diagnosis of each study participant is represented in the x-axis, blue represents control participants and red represents untreated Relapsing Remitting Multiple Sclerosis patients (RRMS). The distribution of female (light blue) or male (purple) study participants is represented in the x-axis. The distribution of study participant’s self-reported ancestries (SRA) including AA (burgundy), Hispanics (green), and NHW (Black) represented in the x-axis. Statistical analysis is represented in the heatmap and represents cell cluster association with: age at the participant level (Age p) FDR values ranging from 0.01 (orange) to −0.01 (purple), the cell level (Age c) FDR values ranging from 1.02 (orange) to 0.98 (purple); with sex at the participant level (Sex p) FDR values ranging from 0.2 (orange) to 0.6 (purple) and the cell level (Sex c) FDR values ranging from 1.4 (orange) to 0.6 (purple) and with **genetic ancestry** PC1 p and PC2 p at the participant level FDR values ranging from 3 (orange) to −2 (purple) and the cell level PC1 c, and PC2 c, FDR values ranging from 1000 (dark orange) to 200 (light orange). * Marks statistically significant associations with FDR <0.05. (**C** to **E**) Correlation of different subpopulations of immunosenescent T cell clusters co-expressing KLRG1 and projected at the cell level.

**Supplementary Figure 3:**
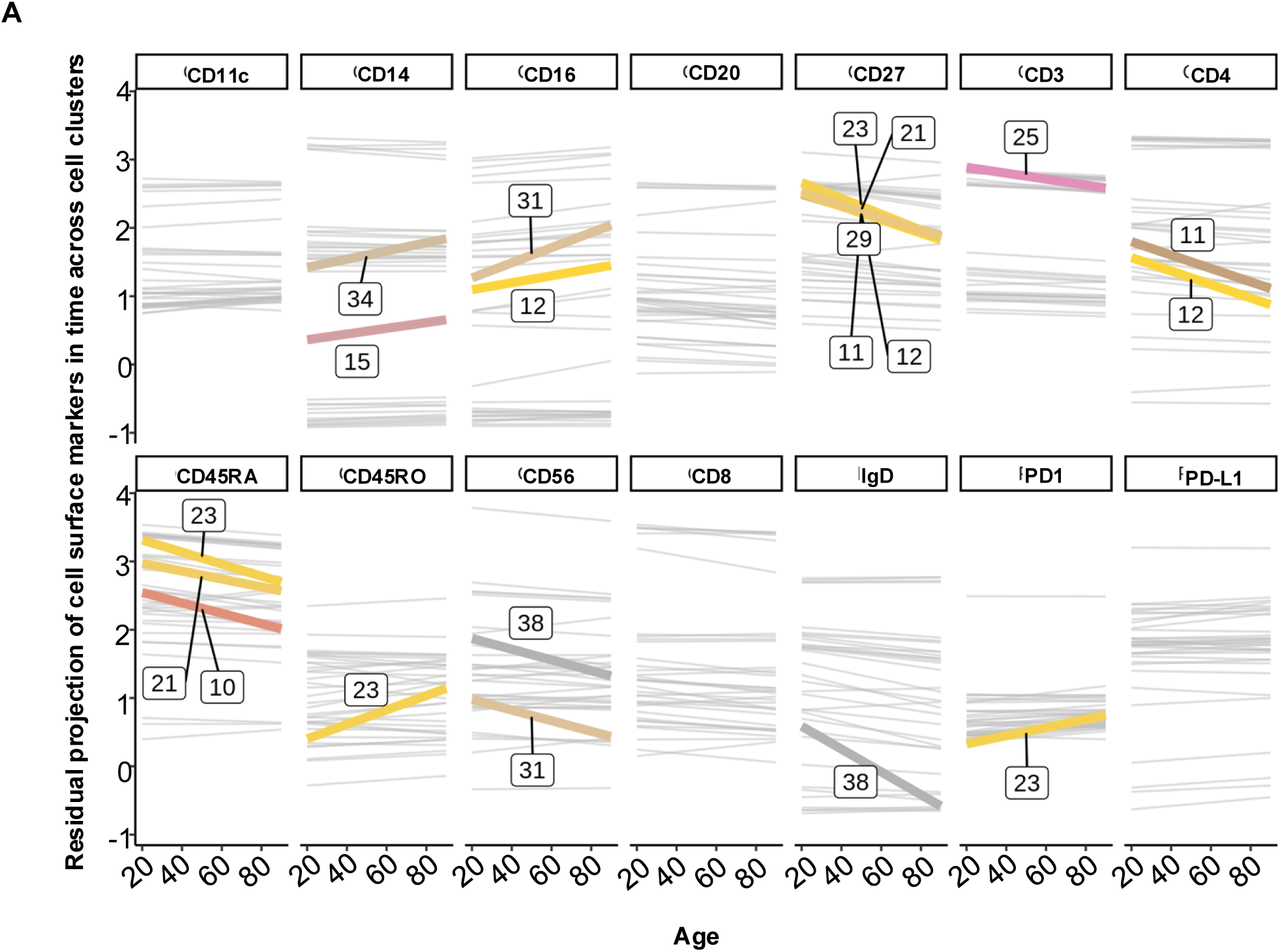
Projecting single cell surface markers within each cell cluster and across the age span. The automated analysis approach enables the evaluation of the expression of a single cell surface marker within each cell cluster. Clusters here are labeled from 1-39. Each colored line defines trajectory of single cell surface markers in association with chronologic age, and each cell surface marker is labeled on top of each regression model plots.

**Supplementary Figure 4:**
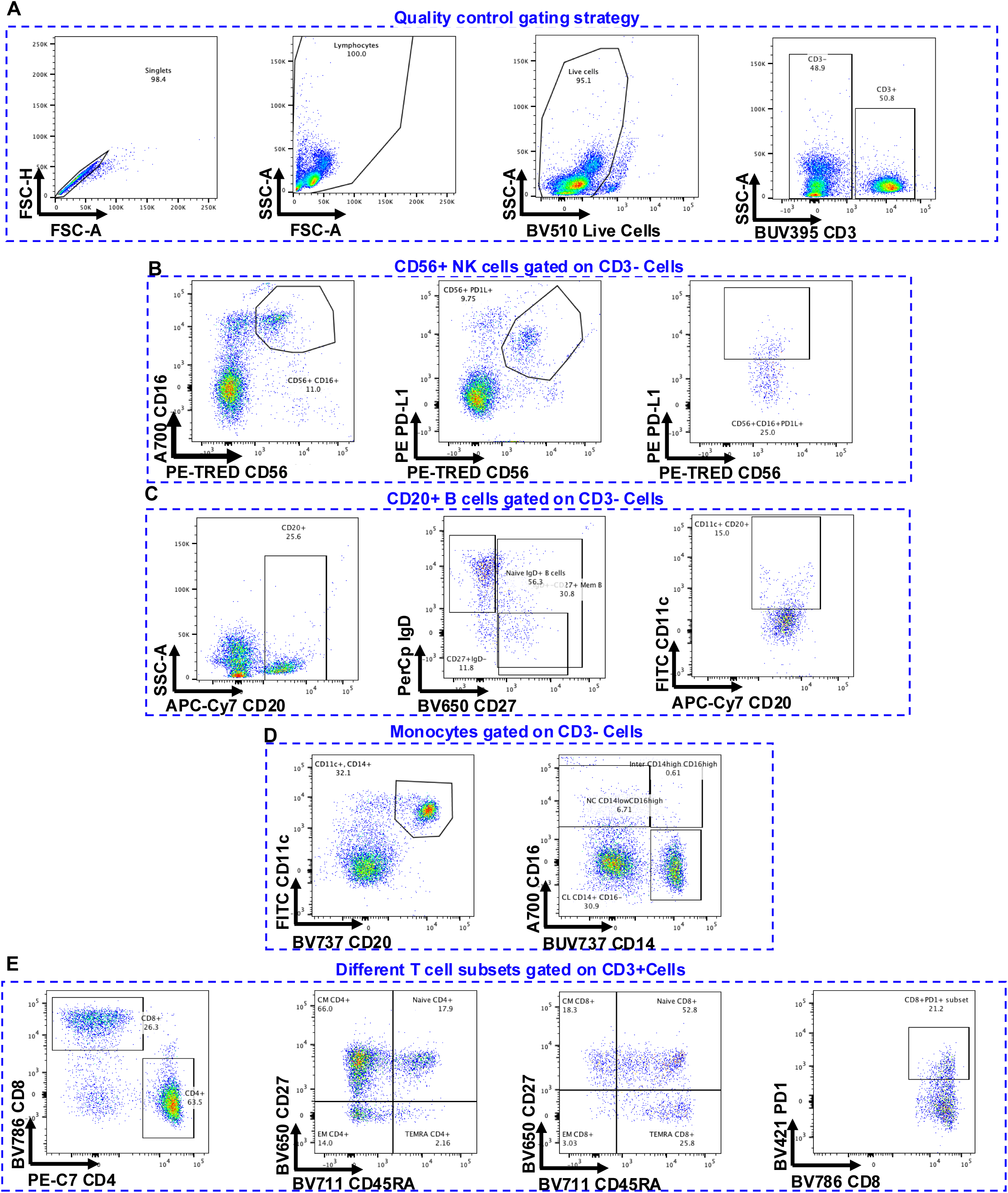
Gating strategy to capture immune cell subsets. relevant to the age associate analysis. (**A**) Gating strategy demonstrating the quality control steps undertaken to gate of defined lineage immune cell subsets. (**B**) Gating on CD3^-^ lymphocytes, further gated on mature CD16^+^ CD56^+^ NK cells, on CD56^+^ PD-L1^+^ NK cells and on CD16^+^ CD56^+^ PD-L1^+^ NK cells. (**C**) Gating on CD3^-^ lymphocytes, further gated on CD20^+^ B cells, on CD27^+^ memory B cells, and CD27^-^ IgD^+^ naïve B cells and on age associated CD11c^+^ CD20^+^ B cells. (**D**) Gating on CD3^-^ lymphocytes, further gated on CD11c^+^ CD14^+^ monocytes, and on CD14 and CD16 to select the non-classical CD14^Low^ CD16^High^ monocytes, the intermediate CD14^High^ CD16^High^ monocytes and theCD14^+^ CD16^-^ classic monocytes. (**E**) Gating on CD3^+^ lymphocytes, on either CD8 or CD4 T cells to captures CD27^+^ CD45RA^+^ Naïve, CD27^-^ CD45RA^+^ TEMRA, CD27^-^ CD45RA^-^ Effector Memory (EM) CD27^+^ CD45RA^-^ Central Memory (CM) T cells, or CD8 T cells co-expressing PD1.

**Supplementary Figure 5:**
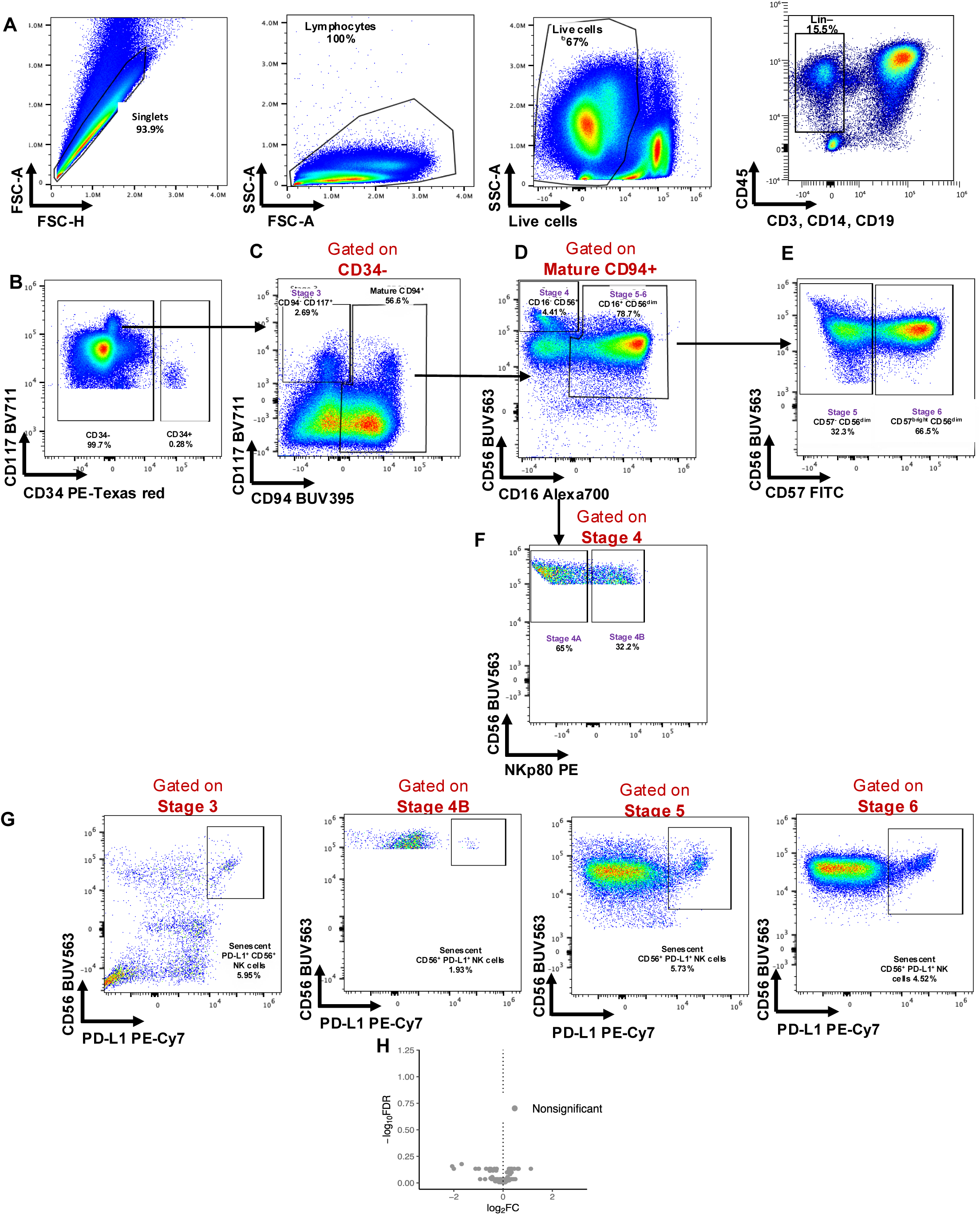
Representative Flow Cytometry gating strategy characterizing the distribution of senescent NK cell subsets across the different maturation stages NK cells (representative plots of an 84 years old participant) Thawed PBMCs were labeled with Live/Dead Stain and further stained with xx antibodies as detailed in the reagent **Supplementary Table 1**. (**A**) Cells were first gated singlets (FSC-A vs. FSC-H), then gated on FSC-A vs. FSC-H to exclude debris. Live cells were identified by gating on Live Dead Stain negative cells. Gating out CD45^+^ lymphocytes as well as CD3^+^, CD14^+^ and CD19^+^ cells, helps to gate on innate immune cells. (**B**) Immature NK cells are defined by CD34^-^. (**C**) Maturation stage 3 NK cells are defined by CD34^-^ CD94^-^ CD117^+^ co-expression, and mature cells are defined by CD94^+^. (**D**) Maturation stage 4 NK cells are defined by CD16^-^ and CD56^+^. (**E**) Maturation stage 5 NK cells are defined by CD57^-^ CD56^Dim^ co-expression, and maturation stage 6 NK cells are defined by CD57^bright^ CD56^Dim^ co-expression. (**F**) NK cell stage 4A and 4B. (**G**) Represent percentages of age-associated CD56^+^ PD-L1^+^ NK cells within each maturation stage, which started to appear at stage 4B. (**H**) Differential gene expression (DEG) comparing cluster 13 and cluster 2 using CITE-Seq data. No significant differences in DEG comparing cluster 13 and cluster 2.

**Supplementary Figure 6:**
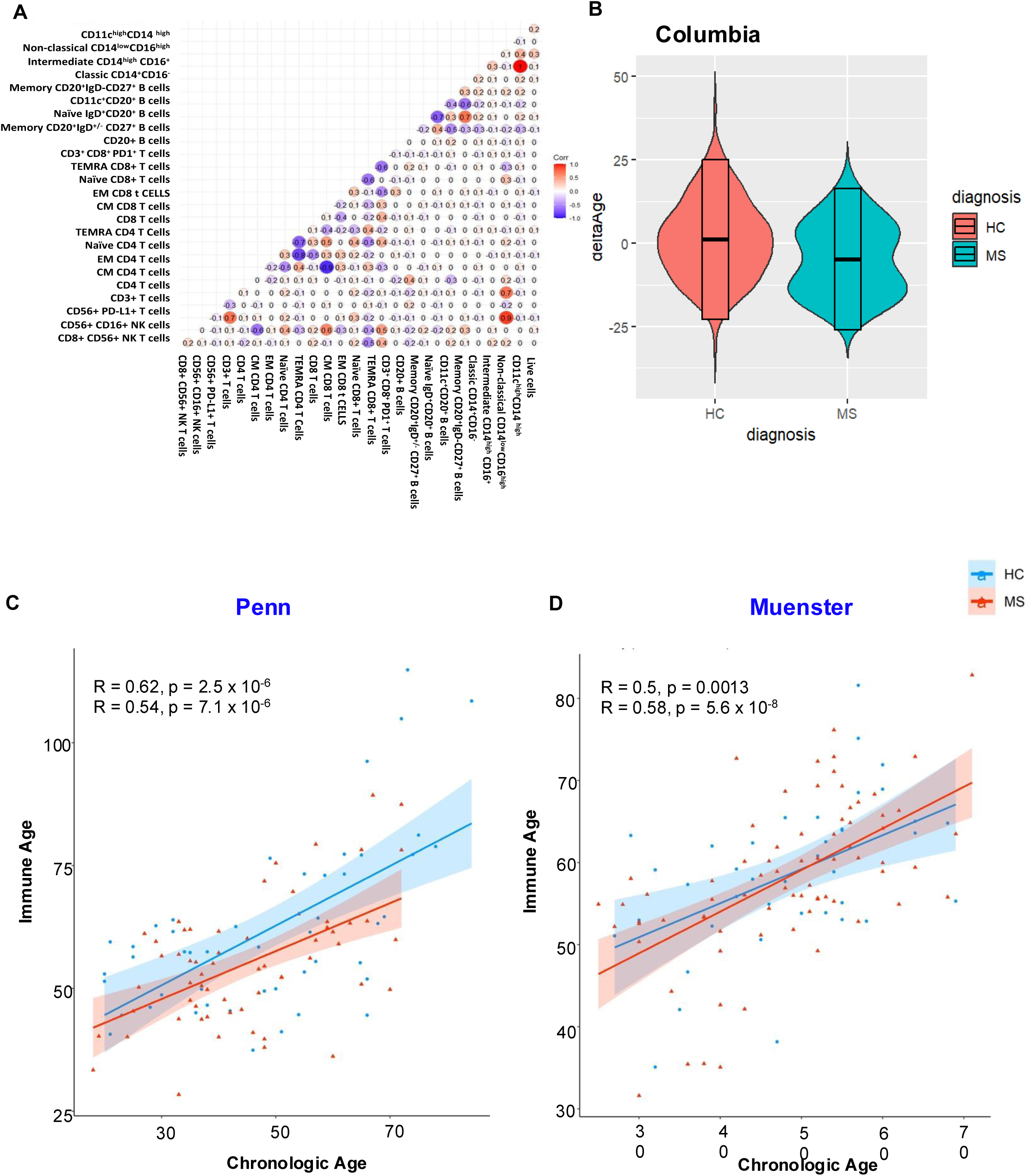
Comparing immune age estimates of control and Multiple Sclerosis (MS) groups. **(A)** Correlation Matrix of frequencies of immune cell subsets curated using the manual FlowJo analysis, using elastic net regression to select best feature of immune age we considered 11 immune cell subsets based on leave-one out cross-validation model. Correlations are attributed scores (+1 to −1) indicating positive *vs.* negative correlations. Delta age was calculated considering immune age estimates and chronologic age. (**B**) Violin plots were used to represent delta age differences comparing control participants with MS patients amongst the Columbia cohort. (**C**) Correlation of immune age and chronologic age using elastic net regression model within the Penn dataset. (**D**) Correlation of immune age and chronologic age using elastic net regression model within the Muenster dataset. Each blue dot represents a control participant, and each red dot represents an MS patient.

**Supplementary Figure 7:**
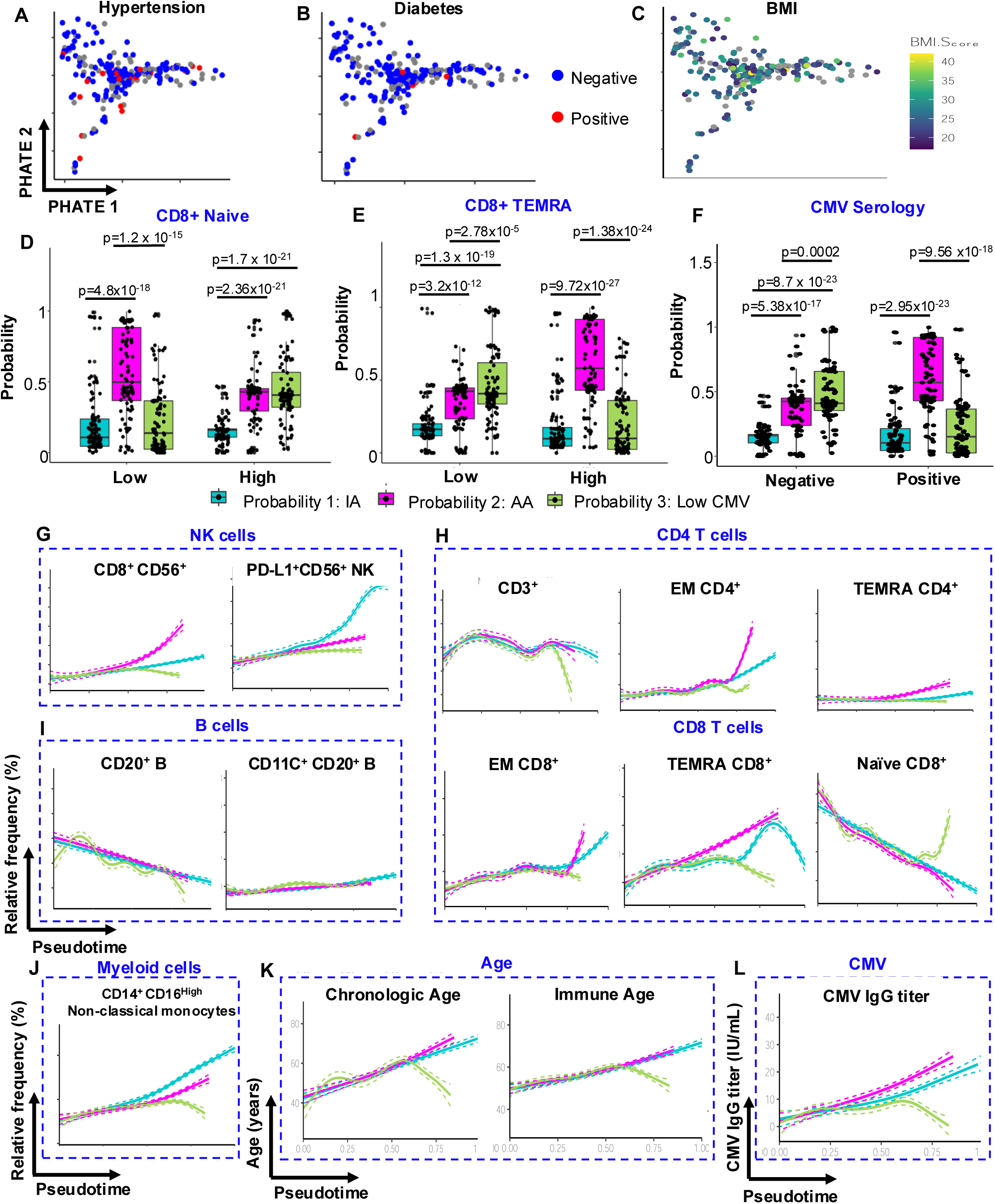
Trajectories of immunosenescence. PHATE Representation of distinct patterns of immune aging trajectories correlated with (**A**) participant’s hypertension status, (**D**) participant’s diabetes status, (**C**) participant’s body mass index (BMI). (**D**) Representative plots highlighting the distribution of the three different probabilities within “low” *vs.* “high” CD27^+^ CD45RA^+^ naïve CD8^+^ T cells participants. Participants were grouped as “low” if frequencies of CD27^+^ CD45RA^+^ naïve CD8^+^ T cells were < 34%, or as “high” if frequencies of CD27^+^ CD45RA^+^ naïve CD8^+^ T cells were > 34 %. (**E**) Representative plots highlighting the distribution of the three different probabilities within “low” *vs.* “high” CD27^-^ CD45RA^+^ TEMRA CD8^+^ T cells participants. Participants were grouped as “low” if frequencies of CD27^-^ CD45RA^+^ TEMRA CD8^+^ T cells were < 21.4%, or as “high” if frequencies of CD27^-^ CD45RA^+^ TEMRA CD8^+^ T cells were > 21.4%. (**F**) Representative plots highlighting the distribution of the three different probabilities within CMV seronegative *vs.* CMV seropositive participants group. (**G** to **J**) Represents the projection of pseudotime with relative frequencies of immune cell subsets that are significantly changing with age, and projection across the three different probabilities. Each dot represents cell subset frequencies derived from individual study participants. (**G**) NK cells, (**H**) CD4 and CD8 T cells, (**I**) B cells, and (**J**) Myeloid cells. (**K)** Represents the projection of pseudotime correlated with chronologic and immune age, and across the three different probabilities. (**L**) Represents the projection of pseudotime correlated with CMV IgG titers across the three different probabilities. Each dot represents a data point from individual donors.

