## Supplementary Table 2 for "Relation of CMV and brain atrophy to trajectories of immunosenescence in diverse populations"

| **Marker** | **Clone** | **Fluorochrome** | **Catalogue #** | **Source** |
| --- | --- | --- | --- | --- |
| CD3 | SK7 | BUV395 | 564001 | BD Bioscience |
| CD4 | SK3 | PE-Cy7 | 557852 | BD Bioscience |
| CD8 | RPA-T8 | BV786 | 563823 | BD Bioscience |
| CD11c | B-ly6 | FITC | 561355 | BD Bioscience |
| CD14 | M5E2 | BUV737 | 612763 | BD Bioscience |
| CD16 | 3G8 | Alexa Fluor 700 | 302025 | BD Bioscience |
| CD19 | 2A3 | BUV496 | 612938 | BD Bioscience |
| CD20 | 2H7 | APC-H7 | 56853 | BD Bioscience |
| CD27 | M-T271 | BV650 | 564894 | BD Bioscience |
| CD28 | CD28.2 | BV421 | 562613 | BD Bioscience |
| CD34 | 581 | PE-Dazzle 594 | 343533 | Biolegend |
| CD45 | Hi30 | BUV805 | 612891 | BD Bioscience |
| CD45RA | HI100 | BV711 | 563733 | BD Bioscience |
| CD45RO | UCHL1 | APC | 5611137 | BD Bioscience |
| CD56 | HCD56 | PE-Dazzle 594 | 318348 | Biolegend |
| CD57 | NK-1 | PE-Dazzle 594 | 562488 | BD Bioscience |
| CD94 | HP-3D9 | BUV394 |  |  |
| CD95 |  |  |  |  |
| CD103 | Ber-ACT8 | BV785 | 350229 | Biolegend |
| CD107a | H4A3 | V450 | 561345 | BD Bioscience |
| CD117 | 104D2 | BV711 | 313229 | Biolegend |
| CD159c (NKG2C) | 134591 (RUO) | BV480 | 748168 | BD Bioscience |
| CD159a (NKG2A) | 131411 | BUV615 | 226946 | BD Bioscience |
| CD294 | BM16 | BV510 | 350119 | Biolegend |
| CCR7 | 2-L1-A |  |  |  |
| Granzym B | QA16A02 | PerCP/Cy5.5 | 372211 | Biolegend |
| IFN-g | 4S.B3 | BV650 | 502537 | Biolegend |
| IL-10 | JES3-19F1 | PE | 506801 | Biolegend |
| IgD | IA6-2 | BUV615 | 613008 | BD Bioscience |
| Ki67 | BUV737 | B56 | 567130 | BD Bioscience |
| KLRG-1 (MAFA) | SA231A2 | AF647 | 51-5893-82 | ThermoFisher |
| CD336 (NKP44) | p44-8 | BV605 | 474301 | BD Bioscience |
| Perforin | dG9 | BV421 | 308121 | Biolegend |
| PD1 | EH12.2H7 |  |  |  |
| PD-L1 | MIH3 | PE/Cy7 | 374505 | Biolegend |
